# RUBRIC: rapid quality assessment tool for RNA 3D structures

**DOI:** 10.64898/2026.11.20.767675

**Authors:** Kamshinen J. Dewan, Joseph D. Yesselman

## Abstract

Cryo-electron microscopy is revealing increasingly complex RNA structures, but deposited models can contain distorted base pairs, incorrect ribose puckers and backbone conformational outliers. Here, we introduce Rapid Unified Base-pair, Ribose and backbone Integrity Check (RUBRIC), an open-source program for integrated RNA annotation and geometric validation. RUBRIC classifies base pairs using the Leontis-Westhof scheme, flags ambiguous assignments and scores unambiguous pairs against reference geometries, identifying the parameters responsible for low scores. It also assigns backbone suites to known conformers, proposes target torsions for unassigned suites and flags incorrect ribose puckers. RUBRIC analyzes a complete human ribosome in under one second, with base-pair annotations consistent with leading programs. Disagreement among those programs was more frequent for pairs flagged as ambiguous, supporting RUBRIC’s identification of uncertain classifications. Flagged geometric abnormalities were reduced in higher-resolution models of the same RNA or following targeted re-refinement, supporting their relevance to correctable modeling problems. A PyMOL plug-in and machine-readable outputs support model inspection and correction. RUBRIC enables rapid, interpretable assessment of RNA geometry, providing a foundation for improved structural models and better datasets for training and evaluating RNA structure prediction methods.

## Introduction

RNA molecules carry out diverse biological functions through three-dimensional (3D) folds stabilized by base pairing and, in many RNAs, tertiary contacts [1–6]. Canonical Watson-Crick pairs predominate in helices, whereas noncanonical pairs help organize loops, junctions and long-range interactions [7–9]. Advances in X-ray crystallography and especially cryo-electron microscopy (cryo-EM) have rapidly expanded the number and complexity of experimentally determined RNA structures [3,4,10,11]. Many cryo-EM maps are determined at moderate resolution and may not be able to distinguish between alternative local geometries. This can allow models with distorted base pairs, incorrect ribose puckers or backbone errors to fit the density [12–14]. The increasing use of these structures in mechanistic studies and in training and evaluating structure-prediction methods creates a pressing need for rapid geometric validation to identify potential modeling errors before they propagate into downstream analyses [8,15–17].

During deposition to the Protein Data Bank (PDB), the wwPDB validation pipeline assesses agreement between the atomic model and experimental data and evaluates model geometry [18]. For RNA, geometric checks include bond lengths and angles, steric clashes, backbone conformations and ribose puckers [19,20]. Backbone and ribose-pucker outliers are flagged for depositor review but do not automatically prevent deposition; inclusion in the PDB therefore does not establish that a model is free from significant local conformational errors [18]. Standard validation reports also lack a comprehensive quantitative assessment of canonical and noncanonical base-pair geometry, despite the central role of these interactions in organizing RNA secondary and tertiary structure [7,21]. Furthermore, refinement restraints can maintain acceptable covalent geometry without ensuring that interacting nucleotides are correctly positioned [18,22].

Several complementary approaches address these problems through rebuilding, re-refinement and structural analysis. RNA-specific methods, including RNABC, RCrane and ERRASER, can correct backbone conformations, sugar puckers and steric clashes through conformational sampling and, where available, guidance from experimental density [23–25]. PDB-REDO and CERES automate the re-refinement of deposited crystallographic and cryo-EM models, respectively [21,26]. Force-field-based refinement and molecular-dynamics approaches combine physical models with experimental observations, including ensemble refinement to account for RNA conformational heterogeneity [27–29]. Other methods characterize noncanonical interactions and quantify their geometric similarity to reference structures [30–32], while DNATCO integrates backbone and covalent-geometry validation with base-pair annotation [33]. Building on these advances, our ultimate goal is to incorporate quantitative assessment of RNA interactions directly into refinement software, where it could help identify local geometric problems and guide their correction during coordinate optimization bypassing the need to perform re-refinement. The present study takes a first step toward this goal by developing a rapid, interpretable scoring framework that assesses base-pair geometry alongside backbone conformation and ribose pucker. This framework provides a foundation for future integration into refinement workflows and for systematic evaluation of structural datasets used in downstream AI applications.

Base pairs are the principal elements of RNA secondary and tertiary structure. The Leontis-Westhof (LW) scheme defines twelve base-pair classes according to the interacting edges of the two bases, Watson-Crick, Hoogsteen or sugar, and the cis or trans orientation of their glycosidic bonds [7,34]. Established programs identify base pairs and assign these classes. However, two limitations remain. First, uncertainty in classification is not consistently communicated. While the Leontis-Westhof system has transformed how base pairing is classified, base-pair geometry is continuous and does not always fit into Leontis-Westhof classes. Interactions near class boundaries can receive different assignments from different programs [8,31,35,36]. Some tools mark near pairs or undetermined edges, but do not necessarily identify the competing classes, leaving users to interpret uncertain assignments [32,37,38]. Second, annotation programs do not provide an overall assessment of base-pair geometric quality. ClaRNA [31] comes closest, scoring how closely each pair matches reference examples of its class and reporting intermediate pairs as imperfect matches to two classes, but this single score does not show whether a low value reflects an intermediate geometry or a distorted one. A pair can be confidently identified and classified yet have distorted relative base orientations or unfavorable hydrogen-bond geometry. Thus, its class label alone does not reveal these problems or indicate their severity. Base-pair analysis therefore needs to report annotation uncertainty and geometric quality as separate properties, making clear both how confidently an interaction is classified and how well its geometry conforms to appropriate reference structures.

To address these gaps, we developed RUBRIC (Rapid Unified Base-pair, Ribose and backbone Integrity Check), an open-source program that combines base-pair annotation and geometric quality assessment with checks of RNA backbone conformation and ribose pucker. RUBRIC explicitly reports classification uncertainty, identifying both candidate Leontis-Westhof classes when a pair’s geometry does not clearly favor either. Its quality scores assess unambiguous pairs against reference geometries and identify the parameters responsible for low scores. The program also evaluates backbone suites against 54 reference conformers and flags ribose-pucker inconsistencies [12,20]. RUBRIC analyzes a complete human ribosome in under a second, enabling repeated assessment during refinement. Its annotations agree with those of established programs, and pairs flagged as ambiguous are more often classified inconsistently by other tools. Flagged geometric problems generally improve in higher-resolution models, while PDB-REDO re-refinement improves primarily the Watson-Crick geometry it restrains. These results support RUBRIC’s ability to identify correctable modeling problems. Its local, machine-readable assessments can guide model correction and curation of RNA structural datasets.

## Methods

### Overview and implementation

RUBRIC is a stand-alone open-source program (https://github.com/YesselmanLab/RUBRIC, installable with pip install rubric-rna) that takes a single RNA-containing coordinate file (mmCIF or PDB format, optionally gzip-compressed) and, in one pass, (i) identifies base pairs, (ii) assigns each pair a Leontis-Westhof (LW) geometric class [7], (iii) scores the local geometric quality of every base pair with a single LW class against empirical distributions derived from curated reference structures, and (iv) assesses every backbone suite against known conformers [20] and assesses ribose pucker [12] (**Figure 1**). The workflow follows the general architecture of established annotation programs, in which a standard base reference frame [39] is fitted to every nucleotide and base pairs are identified from the relative position and orientation of those frames and from hydrogen bonding [32,38,40]. It adds explicit reporting of geometrically ambiguous classifications and a per-interaction quality score. Each stage is described below in the order in which it is executed.

**Figure 1:**
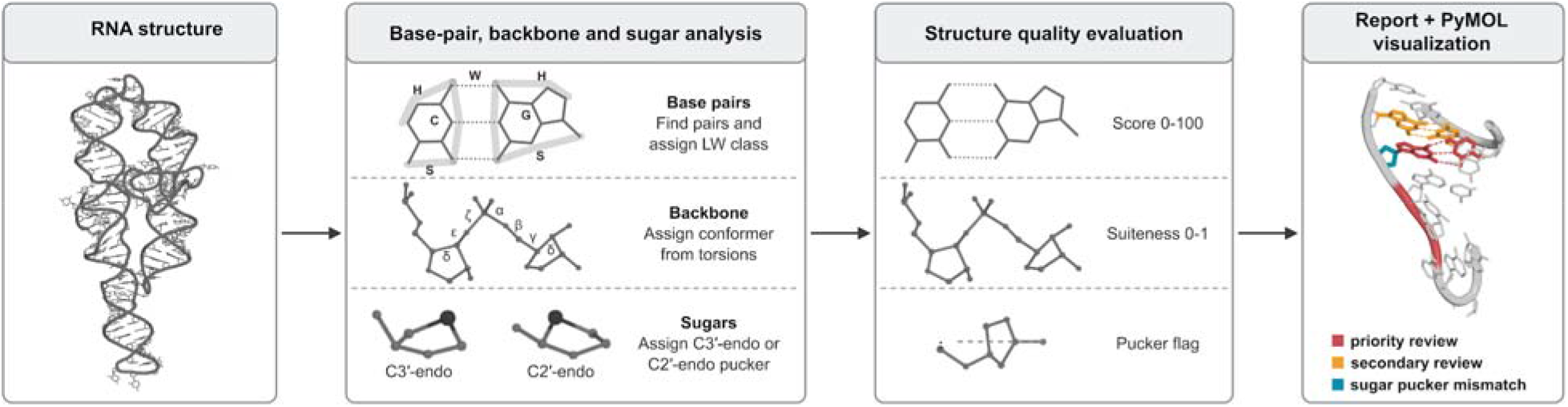
Overview of the RUBRIC pipeline. A single mmCIF or PDB file is analyzed in one pass: base pairs are found and classified, backbone suites are assigned to conformers and ribose puckers are assigned, and each is then evaluated for quality. In the PyMOL view, base pairs in priority review are red, those in secondary review yellow to orange, and riboses with a pucker outlier cyan.

RUBRIC is written in modern C++20 and reads coordinates with GEMMI (v0.6.5) [41]. All reference data are distributed with the program, so no external annotation program or database is required, and local files are analyzed without network access. For each structure, RUBRIC writes a JSON report of the selected base pairs with their LW classes, hydrogen bonds, rigid-body parameters, scores, review tiers and itemized penalties, together with structure-level summaries and per-residue backbone results. A complete analysis is run with a single command (`rubric model.cif`); all options are listed in **Supplementary Table S1**.

### Base reference frames

Each residue is compared with the standard geometry of its base, the idealized base coordinates defined in the standard reference frame [39] and distributed with the open-source 3DNA program (v2.4) [40,42]. A residue is accepted as a nucleotide if its base-ring atoms superimpose onto the standard base ring with a root-mean-square deviation (RMSD) of at most 0.2618 Å, the 3DNA default. The reference frame of the residue is then obtained by least-squares superposition of the standard base onto the residue’s ring atoms, and modified nucleotides use the standard base of their parent. All superpositions use the closed-form quaternion method [43].

### Identification of candidate base pairs

Following 3DNA [40,42], every pair of residues whose frame origins lie within 15 Å is examined, and a candidate is retained if the vertical separation of the bases is at most 2.5 Å and the angle between the base normals is at most 70°, relaxed from the 65° of 3DNA to retain strongly buckled and propeller-twisted pairs. The glycosidic N1/N9 atoms must also be at least 4.5 Å apart, as in the 3DNA program (v2.4). Sequence neighbors in the same chain (|*i* − *j*| = 1) are not considered. Stacked bases can also pass these limits, so a candidate is removed when one ring center lies more than 2.8 Å above the other and further above it than to the side.

### Hydrogen-bond identification

Donor and acceptor atoms are taken from built-in tables for the standard nucleotides and amino acids, extended to modified residues with a residue chemistry database built from the PDB Chemical Component Dictionary [44]. For each candidate pair, all donor-acceptor contacts within 4.0 Å are collected. Directionality is evaluated by predicting the positions of donor hydrogens and acceptor lone pairs from the local covalent geometry and scoring how well they point along the donor-acceptor vector. Contacts are accepted as hydrogen bonds when the donor hydrogen or acceptor lone pair points along the donor-acceptor vector. Each hydrogen or lone pair can take part in only one bond. Only pairs with at least one base-base hydrogen bond are classified. A pair is selected and scored only if at least one of its base-base hydrogen bonds is 3.5 Å or shorter, whereas its other bonds may be up to 4.0 Å long, as in pairs with possibly distorted geometry, which are therefore still captured and scored.

### Leontis-Westhof classification

Each pair is assigned to one of the 12 Leontis-Westhof classes [7] from the relative orientation of its glycosidic bonds and the edge used by each base. Following RNAView [38], a pair is cis when the two glycosidic bond vectors, projected onto the plane perpendicular to the line joining the base-ring centroids, point to the same side, and trans otherwise. The edge of each base (Watson-Crick, W; Hoogsteen, H; or sugar, S) is first assigned from the direction of the partner base relative to the centroid of each edge’s atoms, and then refined with hydrogen-bond evidence: a bond to an atom that belongs to only one edge fixes that edge, and close contacts with edge-specific atoms resolve the remaining W/H and W/S cases. Each class is named by the orientation (c, cis; t, trans) followed by the edges of the two bases, so that the canonical Watson-Crick pair is cWW.

When both bases are hydrogen-bonded only through atoms shared by two edges and the geometric evidence does not clearly favor one of them, RUBRIC compares the pair with the reference templates of the two candidate classes. The pair is assigned to the better-fitting class when the two template RMSDs differ by more than 0.5 Å. Otherwise RUBRIC reports both candidate classes as ‘X|Y’ instead of forcing a single assignment. We selected these templates from the curated reference structures: for each base-pair type and LW class, the template is the pair whose geometry is most typical of the pairs that the established annotation programs agree on, the pair closest to their average geometry by RMSD over the base atoms.

### Selection of a consistent set of base pairs

RUBRIC selects a consistent set of pairs, allowing each base edge to participate in at most one selected pair. cWW A-U, G-C and G-U pairs that stack consecutively along the chain are first accepted as helical segments, longest first. Among the remaining candidates, RUBRIC then chooses the non-conflicting set that best matches the reference templates and expected hydrogen bonds. Selection never changes a pair’s LW class, and an ambiguous pair keeps its ‘X|Y’ label.

### Base-pair geometric parameters

For each selected pair, the six rigid-body parameters (shear, stretch, stagger, buckle, propeller and opening) are computed from the two base frames with the 3DNA algorithm [40,42] in the Tsukuba convention [39]. For each base-base hydrogen bond, the donor-acceptor distance and the angles at the donor and the acceptor are recorded.

### Reference structure sets

Unless stated otherwise, analyses used the BGSU non-redundant RNA set [45], which groups redundant structures of the same RNA into equivalence classes and represents each class by one entry. The representative entries at a resolution cutoff of 3.5 Å were used (release 4.21; 2,696 PDB entries). Experimental method, resolution, *R*_work_, *R*_free_, clashscore, bond-length and bond-angle root-mean-square *Z*-scores (RMSZ), RNA suiteness and, for cryo-electron microscopy (cryo-EM) entries, the average Q-score [46] were obtained from the wwPDB validation reports through the RCSB PDB [47]. A curated reference set was defined by the criteria in **Supplementary Table S2**, which combine standard indicators of model accuracy and over-fitting [48–50].

The curated set contains 1,323 entries (955 X-ray and 368 cryo-EM; 878,325 nucleotides). The remaining 1,373 entries form the non-curated set. For every scorable pair (a base-base pair with a single LW class), one record was extracted for the pair, one for each of its base-base hydrogen bonds and one for the backbone torsions of each of its residues. The curated set gave 371,960 pairs, 889,864 hydrogen bonds and 739,658 torsion records; the non-curated set gave 512,247 pairs, 1,228,826 hydrogen bonds and 1,017,973 torsion records.

### Empirical scoring tables

#### Reference groups and structure weighting

All scoring tables were built from the curated set alone. Each parameter of a pair is judged against pairs of the same base-pair type and Leontis-Westhof class (a reference group, for example G-C/cWW). When a group has fewer than ten observations, it is expanded to include all base-pair types within the same LW class, and if the expanded group still contains fewer than ten observations, all pairs are used, regardless of base-pair type or LW class. Each observation is weighted by 1/*n*, where *n* is the number of observations from the same entry in that group, so that large assemblies do not dominate, as in Černý et al. [22]. Tables were built for the six rigid-body parameters, hydrogen-bond distance and donor and acceptor angles.

#### Probability percentile score (ProSco)

The probability percentile score (ProSco) was introduced by Černý et al. [22] to validate the valence geometry of nucleic acids, the bond lengths and angles within each nucleotide, and is used for this purpose in DNATCO [33]. Here we apply it to the geometry of base pairs. ProSco expresses how typical a value is within its group on a 0-100 scale. For each group and parameter, the reference density is estimated with a Gaussian kernel density estimate computed with SciPy (v1.17.1) [51], with each observation weighted by the inverse of the number of observations from its PDB entry, so that large assemblies such as ribosomes, which contribute many observations, do not dominate the reference. The bandwidth, which sets how widely each observation is smoothed, was chosen by unbiased cross-validation and multiplied by 1.5, following Černý et al. Where the cross-validation optimum lay at the edge of its search range, Scott’s rule [52] was used instead. The density is evaluated on equal bins about half a bandwidth wide, and the bins are ranked from highest to lowest density. The ProSco of a value *x* is

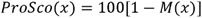

where *M*(*x*) is the fraction of the reference distribution in bins that are denser than the bin containing *x*. A value in the densest bin has a ProSco of 100, and values with ProSco < 5 lie in the least typical regions, which together hold 5% of the reference.

#### Asymmetric non-parametric standard score (Z′)

The distance of a value *x* from the center of its group is measured by the weighted asymmetric non-parametric standard score of Černý *et al.* [22]:

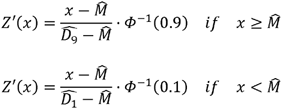

where M□ is the weighted median, D□_1_ and D□_9_ are the weighted first and ninth deciles and Φ⁻¹ is the probit function, so that *Z*′ reduces to an ordinary *z*-score for normally distributed data.

### Base-pair quality score

#### Severity of each parameter

Ten parameters are evaluated for each scorable pair: shear, stretch, stagger, buckle, propeller, opening, hydrogen-bond distance, hydrogen-bond angles, hydrogen-bond count and steric clash. Each receives a severity *s* between 0 (normal) and 1. Following the PDB-based criterion of Černý *et al.* [22], a value is considered usual (Preferred) if its ProSco is at least 5. Where Černý *et al*. label values outside this range as Allowed or Of Concern, we instead penalize each such value with a severity, a number that rises gradually as the value becomes more unusual, so that penalties of the ten parameters can be combined into one score. For the eight continuous parameters, the severity is

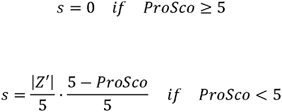

where |Z′|/5 is capped at 1, which corresponds to |Z′| = 5, the threshold that Černý *et al.* [22] use for values of concern. As a result, a value is penalized only if it is as rare as the least typical 5% of the curated reference, the penalty grows with distance from the median, and the factor (5 − ProSco)/5 makes the severity rise continuously from zero at the Preferred boundary.

#### Hydrogen-bond parameters

The hydrogen bonds of a pair are described by three parameters, the hydrogen-bond distance, the hydrogen-bond angles and the hydrogen-bond count. The donor and acceptor angles of every base-base bond are scored against their own tables. For both the distance and the angles, the pair receives the severity of its worst value. The expected number of base-base hydrogen bonds of each group is its most frequent count in the curated set, and a pair with fewer bonds receives a severity equal to the fraction of the expected bonds that are missing.

### Steric clashes

For every pair of heavy atoms of the two residues, the overlap is the minimum acceptable separation minus the observed distance; the minimum is 2.6 Å for atoms that could hydrogen-bond and otherwise the sum of their van der Waals radii. The worst overlap involving at least one base atom contributes to the pair score once it exceeds 0.4 Å, the conventional threshold for a serious clash [16,19,53]; its severity rises linearly from 0 at an overlap of 0.4 Å to 1 at 0.9 Å and stays at 1 for larger overlaps.

### Penalty weights

The penalty weight of each parameter reflects how much greater its mean penalty severity is in structures that do not meet the curation criteria (**Supplementary Table S3**). Every scorable pair of the curated (371,960 pairs) and non-curated (512,247 pairs) sets was scored against the curated tables. For each parameter *i*, the mean severity was computed over pairs in each set, with add-one (Laplace) smoothing to avoid division by zero. The weight is the natural logarithm of the ratio of the two means, set to zero if negative (which did not occur), rescaled so that the ten weights sum to 100:

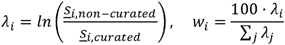

Mean penalty severity is higher for every parameter in the non-curated set (**Supplementary Table S3**), by about twofold for most and threefold for stagger; the hydrogen-bond count is the exception (ratio 1.19).

### Pair score, review tiers and structure-level score

The score of a pair is summed over the ten parameters.

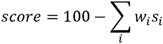

Because the weights sum to 100 and each severity lies between 0 and 1, scores range from 0 to 100, and a pair whose every parameter lies in its Preferred region scores 100. Each penalized parameter is reported as an issue. Pairs are assigned to three review tiers: Preferred (score = 100), secondary review (80 ≤ score < 100) and priority review (score < 80). The cutoff of 80 was chosen because the enrichment of flagged pairs in the non-curated set stayed at about 2.5-fold for cutoffs from 70 to 80 and declined above it, to 2.1-fold at 95 (**Supplementary Figure S1A** and **B**). Priority-review pairs were penalized most often for hydrogen-bond angles (83% of pairs; **Supplementary Figure S1C**). The validation results did not depend on this choice: at cutoffs of 75, 80 and 85, flagged pairs improved in 97-98% of molecules determined at higher resolution and in 87-91% of re-refined structures (**Supplementary Figure S1D**). Ambiguous pairs are reported with a null score. The structure-level base-pair score is the mean score of all scored pairs.

### Backbone conformation and ribose pucker

Backbone torsions α-ζ are computed for suites of residues that are consecutive in the same chain and covalently linked. Each suite is assigned to one of the 54 backbone conformers of the RNA Ontology Consortium [12,20] by a re-implementation of Suitename (https://github.com/rlabduke/suitename) that uses its conformer library and classification parameters unchanged, and receives a suiteness between 0 and 1; unassignable suites (’!!’) have suiteness 0. Suites are assigned only to nucleotides whose C2′ atom is bonded to an O2′ atom, so DNA and nucleotides modelled without the 2′ oxygen receive no suite. The structure’s backbone suiteness is the mean over its scored suites, as in the wwPDB RNA-suiteness metric, which RUBRIC reproduces (R² = 0.992; 2,604 structures; **Supplementary Figure S2A**). For residues with suiteness below 0.5 or an unassignable suite, RUBRIC lists the torsions that are atypical for the conformer (ProSco < 5 in per-conformer tables built from the curated set) and suggests a target value toward the center of the assigned conformer, or of the nearest conformer for an unassignable suite.

Ribose pucker is inferred both from δ, using the ranges of Suitename (C3′-endo 60-105°, C2′-endo 125-165°), and from the perpendicular distance (Pperp) between the 3′ phosphorus and the extended glycosidic bond, with Pperp ≥ 2.9 Å indicating C3′-endo and Pperp < 2.9 Å indicating C2′-endo [12]; residues whose two assignments disagree are reported as pucker outliers (**Supplementary Figure S2D**). C2′-endo riboses in canonical cWW pairs, where the A-form C3′-endo pucker is expected, are flagged separately as unusual puckers [54]. Residues with δ outside both ranges are reported as unclassified, together with the pucker implied by the phosphate position. Because each suite spans two consecutive riboses, δ(i−1) to δ(i), and Suitename sorts both δ values into these two ranges before assigning a conformer, an unclassified δ makes the one or two suites that contain it unassignable (suiteness 0). These flags do not affect any score.

### Interactive visualization in PyMOL

RUBRIC includes a plug-in for PyMOL [55]. A single command (rubric-view, followed by a file name or PDB identifier) loads the structure, runs RUBRIC and displays the results, and the same analysis can be run on any object in an open session. Flagged base pairs are shown as sticks colored by score (red for priority review, yellow to orange for secondary review), and riboses with a pucker outlier are marked with a cyan ring. Flagged pairs form a worklist ordered by tier and severity; the user can center on each entry, list its score and issues and superimpose the reference template of its base-pair type and LW class. Re-running the analysis after the model is edited updates the display. With --pml, RUBRIC also writes a PyMOL script that reproduces the colored display and the worklist, as a named selection for each priority-review pair, so that the results can be viewed without the plug-in (**Supplementary Table S1**). Neither the plug-in nor the script modifies the coordinates. All structure images in the figures were rendered with PyMOL [55].

### Comparison with established annotation tools

RUBRIC was compared with DSSR [32], FR3D [37], RNAView [38], barnaba [56], RNApolis Annotator [57] and BPNet [58]. The six programs were chosen as widely used programs that classify base pairs in the Leontis-Westhof scheme and read mmCIF files directly. All programs were run locally on the same mmCIF files with the versions and commands in **Supplementary Table S4**. With DSSR as the annotation reference, recall before pair selection for each LW class over all entries of the non-redundant set was the fraction of DSSR’s pairs that RUBRIC classified in the same class, whether or not selection kept them, and precision after pair selection was the fraction of RUBRIC’s selected pairs that DSSR assigned to the same class [31,59]. These metrics describe different stages of the RUBRIC pipeline and measure agreement with DSSR. Agreement among all seven programs was measured on the 2,660 entries of the non-redundant set that every program processed. Only RUBRIC’s selected pairs were compared, and ambiguous pairs were excluded. Detection agreement was measured with the Jaccard index [60]. Class agreement of pairs detected by both programs was measured with Cohen’s κ [61]. Detection agreement computed with all pairs that RUBRIC classified, whether selected or not, is given in **Supplementary Table S5**.

Pairs that RUBRIC and DSSR [32] assigned to different classes were compared with the annotations of the other five programs (**Supplementary Figure S3C**). Each of these pairs was also compared with the templates of the RUBRIC class and the DSSR class by RMSD, and differences of 0.1 Å or less were treated as ties (**Supplementary Figure S3D**). To test whether ambiguous pairs are those on which other programs disagree, a RUBRIC pair classified by at least two of the other programs was counted as disputed if they assigned more than one class, and the rate for ambiguous pairs was compared with that for unambiguous pairs of the same classes. Differences were tested with two-sided z-tests with cluster-robust variance over structures.

### Runtime benchmark

Runtime was measured on 2,096 mmCIF files from the non-redundant set (4 to 15,870 nucleotides). Each program was run with its native command line (**Supplementary Table S4**), and wall-clock time was measured from process launch to exit, including file reading and start-up. Each program-structure combination was timed three times in random order, with one thread per process and four structures processed concurrently on an Apple M2 Pro (16 GB memory), and the median time was used. Structures on which a program failed were excluded for that program only. The speed-up was the ratio of the other program’s median time to RUBRIC’s on the same structure (**Supplementary Figure S4G**).

### Validation against independently improved models

To test whether the base pairs, backbone and ribose features that RUBRIC flags are correctable, we compared original and improved models of the same molecule obtained without reference to RUBRIC, from two sources. First, re-refined models of the X-ray entries of the non-redundant set were obtained from PDB-REDO [62] and scored together with the deposited models, so that the experimental data are identical and only the refinement differs. At resolutions worse than 1.7 Å, PDB-REDO restrains base-pair hydrogen bonds of Watson-Crick pairs (detected with DSSR) and base stacking, but applies no restraints to backbone torsions or ribose pucker [21]. Second, molecules deposited more than once were identified from the BGSU equivalence classes. Within each class whose members were all determined by the same experimental method, depositions differing in resolution by at least 0.5 Å were paired, the lower-resolution entry being the original. Pairs were taken greedily in order of decreasing resolution difference, at most one per class and without reusing an entry, and were accepted if the two chains shared at least 30 residue numbers with at least 95% sequence identity at those positions. Depositions were paired on resolution alone, independently of RUBRIC scores, giving 162 molecules (99 cryo-EM and 63 X-ray; resolution differences 0.5-1.9 Å, median 0.76 Å). Base pairs were matched by residue identifier and LW class, within the chosen chains by residue number and base identity at higher resolution, and suites and riboses were matched by residue identifier. The unit of comparison was the structure for re-refinement, in which the original and improved models are the same entry, and the molecule for the comparison at higher resolution, in which they are two entries.

Four measures were computed for each original and improved model over matched objects, namely the mean score of the priority-review pairs of the original model, the mean suiteness, the percentage of unassignable suites and the number of riboses with a pucker outlier, and each unit contributed one value per measure so that large assemblies count once. For each issue type, its frequency among a unit’s priority-review pairs was computed in both models and averaged across units. Flagged pairs were also divided into canonical (A-U and G-C cWW) and noncanonical (all others, including G-U) pairs, to separate corrections of the geometry that PDB-REDO restrains from those it does not (**Supplementary Figure S5A and B)**. Because pairs were selected for low scores, part of their improvement is expected from regression to the mean, so the analysis was repeated in reverse, selecting pairs flagged in the improved model and following them back to the original. Selection raises the change in both directions, whereas a genuine improvement raises the forward change and lowers the reverse change, so a forward change larger than the reverse change indicates improvement beyond that expected from selection. The two changes were compared with a two-sided z-test on the per-unit differences, using units that contained flagged pairs in both models (227 structures after re-refinement and 61 molecules at higher resolution; **Supplementary Figure S5C**). As a further control, score changes were measured in structures in which every pair scored 100 (preferred tier) before re-refinement (**Supplementary Figure S5D**).

### Relationship with global quality measures

Structure-level base-pair scores were compared with four measures from the wwPDB validation reports (resolution, clashscore, R_free_ of X-ray structures, average Q-score of cryo-EM structures), with RUBRIC backbone suiteness and with RNA length (**Supplementary Figure S6**). Base-pair scores and backbone suiteness were also compared across experimental methods (**Supplementary Figure S7**). For the correlations, structures with fewer than five scored pairs or without the measure being compared were excluded. The comparison between experimental methods included every structure with at least one scored pair. For the rates of unassignable suites (**Supplementary Figure S2B and C**), structures of the curated and non-curated reference sets at 3.5 Å resolution or higher with at least twenty suites were included, and entries sharing at least 80% of their base pairs (Jaccard index) were treated as one molecule, so each molecule contributed one rate to each group. Linear relationships were fitted by ordinary least squares, with RNA length on a logarithmic scale, and are reported as *R*². Solution NMR structures, which are absent from the non-redundant set, were selected in the same way. PDB entries determined by solution NMR that contain RNA and no protein or DNA chains were reduced to one entry per BGSU equivalence class (release 4.62), giving 561 structures with at least one scored base pair, each scored on the first model of its NMR ensemble (**Supplementary Figure S7**).

## Results

### RUBRIC rapidly annotates and assesses base pairs, backbone and riboses with a single command

RUBRIC, written in modern C++20, takes a PDB or mmCIF file and analyzes it with a single command (**Figure 1**). It identifies base pairs and assigns Leontis-Westhof (LW) classes, reporting both candidate classes when the geometry does not clearly distinguish between them. Each unambiguous pair receives a geometric quality score from 0 to 100, accompanied by the parameters responsible for any penalties. Backbone analysis reports conformer assignments and identifies deviating torsions, with suggested target values, for poorly fitting or unassigned suites. RUBRIC also flags ribose-pucker inconsistencies. A PyMOL plug-in displays flagged features directly on the structure for inspection during model building and refinement. Local results and structure-level summaries are provided in a single machine-readable JSON report (**Supplementary Sample Output**), supporting automated analysis and the curation of structural datasets for machine-learning applications, where training-data quality is a recognized concern [8,15].

### RUBRIC annotates and scores a complete human ribosome in under one second

We benchmarked the end-to-end runtime of RUBRIC against six established annotation programs, DSSR [32], FR3D [37], RNAView [38], barnaba [56], RNApolis Annotator [57] and BPNet [58], across 2,096 mmCIF files, using each program’s native command (**Supplementary Table S4**; **Supplementary Figure S4A-F**). RUBRIC annotated and scored complete human 80S ribosomes, comprising approximately 5,500-6,000 nucleotides, in 0.58-0.76 s (**Figure 2A**). It was faster than FR3D, barnaba and RNApolis Annotator on all successfully benchmarked structures. Its advantage over DSSR, RNAView and BPNet increased with RNA size, reaching 11- to 24-fold for structures containing more than 5,000 nucleotides (**Figure 2A**; **Supplementary Figure S4G**). These runtimes make repeated assessment of base-pair geometry, backbone conformation and ribose pucker practical during refinement.

**Figure 2:**
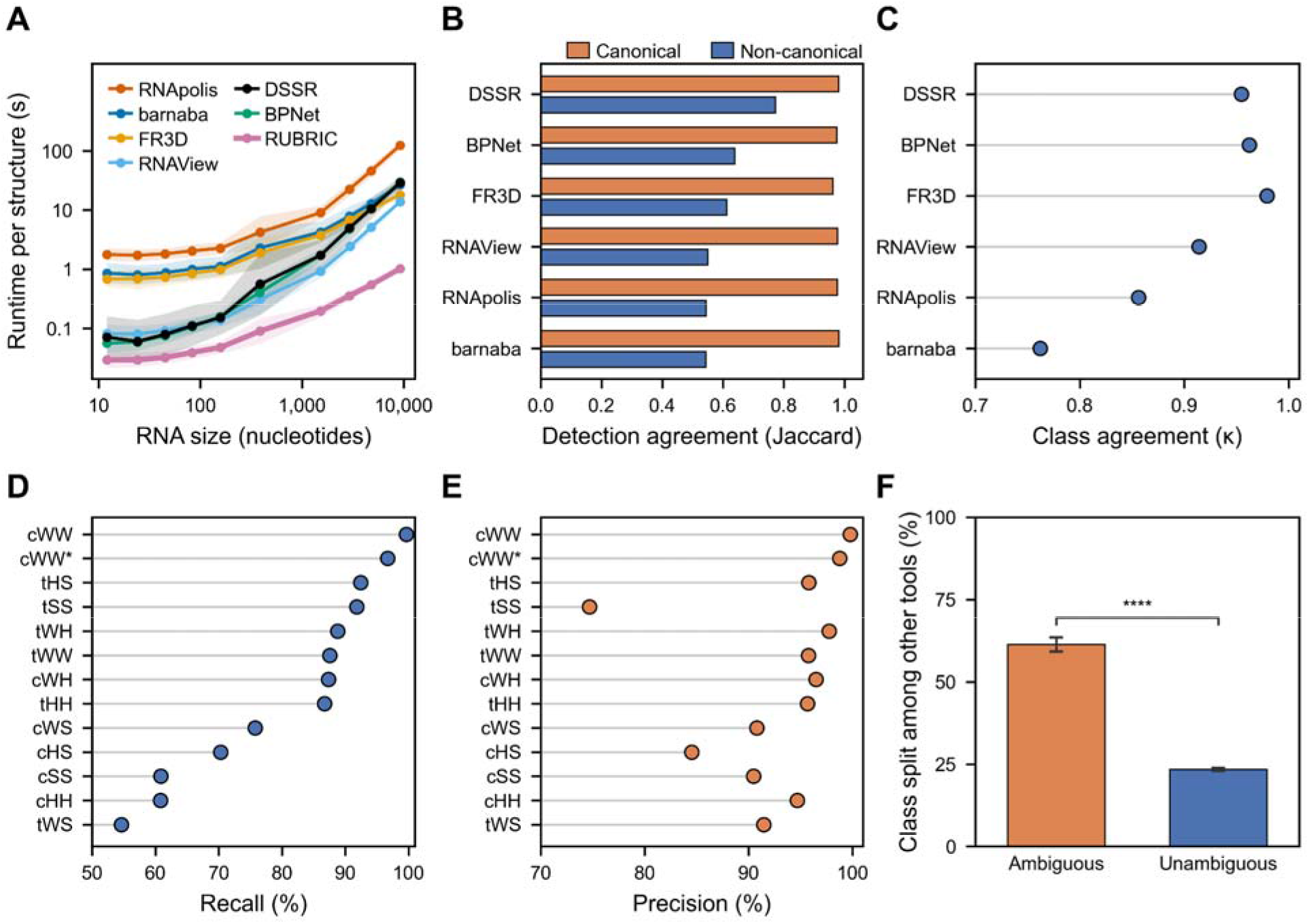
Speed, agreement with six established programs and ambiguity reporting of RUBRIC. (A) Runtime per structure against RNA size (2,045 to 2,096 structures per program). Points are geometric means within logarithmically spaced size bins, merged where needed so that each contains at least 30 structures, and shaded bands span one geometric standard deviation above and below. (B) Agreement in base-pair detection between RUBRIC and each program (Jaccard index) for canonical (orange) and noncanonical (blue) pairs. (C) Agreement in Leontis-Westhof (LW) class between RUBRIC and each program for the noncanonical pairs detected by both (Cohen’s κ). (D and E) Recall before pair selection (D) and precision after pair selection (E) of RUBRIC for each LW class, with DSSR annotations as the reference: recall counts DSSR pairs that RUBRIC classified in the same class, whether selected or not, and precision counts RUBRIC’s selected pairs; cWW* denotes cWW pairs with a noncanonical sequence. (F) Percentage of RUBRIC pairs on which the other six programs assign more than one LW class, for pairs that RUBRIC reports as ambiguous (orange) and for unambiguous pairs of the same classes (blue). Only pairs that at least two of the other programs assign to a single class are counted. Error bars are 95% bootstrap confidence intervals (2,000 resamples of structures); significance is from a two-sided z-test with cluster-robust variance over structures (****, *p* < 0.0001).

### RUBRIC identifies and classifies base pairs consistently with established annotation programs

To assess RUBRIC’s ability to identify and classify base pairs, we compared RUBRIC to six programs listed above on structures from the non-redundant set. We measured detection agreement with the Jaccard index [60] and class agreement among jointly detected pairs with Cohen’s κ [61] (see Methods). RUBRIC showed high detection agreement with each established program for canonical pairs (Jaccard index 0.96-0.98; **Figure 2B**) which is consistent with previous comparisons between annotation programs [31,35]. Agreement was lower for noncanonical pairs: Jaccard indices ranged from 0.42 to 0.70 among established programs and from 0.54 to 0.77 between RUBRIC and those programs (**Figure 2B; Supplementary Figure S3A**). Noncanonical class assignments showed a similar pattern, with κ values of 0.71-0.97 among established programs and 0.76-0.98 for comparisons involving RUBRIC (**Figure 2C; Supplementary Figure S3B**). Only 0.18% of RUBRIC’s noncanonical pairs were reported by no other program, compared with 0.40 to 23.5% for the established programs (**Supplementary Figure S3E**).

To examine class-specific differences, we compared RUBRIC with DSSR, using all RUBRIC-classified candidates to measure recall before pair selection and selected pairs to measure precision after pair selection (see Methods). For canonical cWW pairs, recall was 99.7% and precision 99.8% (**Figure 2D and E**). Across the other eleven classes, both measures were 85-98%, except for recall of tWS pairs (55%), recall of cHH, cSS, cHS, and cWS (61-76%) and precision of tSS pairs (75%). These differences partly reflect different hydrogen-bond requirements. Pairs whose base-base hydrogen bond are all longer than 3.5 are classified by RUBRIC, and therefore count toward recall, but are not selected **(Supplementary Figure S8A).** In addition, RUBRIC requires at least one base-base hydrogen bond, whereas DSSR also accepts pairs whose only hydrogen bond links a base to the partner nucleotide ribose (**Supplementary Figure S8B and C**) [32]. RUBRIC also assigns to tSS some pairs that DSSR assigns to tWS, a classification boundary on which annotation programs are known to disagree [36].

Among the 13,286 pairs detected by both RUBRIC and DSSR but assigned different classes, four of the five remaining programs supported RUBRIC’s assignment significantly more often, by 13-41 percentage points. RNApolis supported the two assignments at similar rates (29% for RUBRIC and 30% for DSSR; **Supplementary Figure S3C**). Comparing each conflicting pair with the templates of its two assigned classes showed that 81.1% were closer to the template corresponding to RUBRIC’s assignment (**Supplementary Figure S3D**). These comparisons support RUBRIC’s consistency with established annotation programs while identifying the interaction definitions and class boundaries responsible for some disagreements.

### RUBRIC flags ambiguous pairs that other programs classify inconsistently

When the geometry of a pair does not distinguish between two LW classes, RUBRIC reports the pair as ambiguous and names both candidate classes instead of forcing one. Pairs flagged as ambiguous by RUBRIC showed greater disagreement among established annotation programs. Among pairs classified by at least two other programs, 61% of ambiguous pairs received conflicting assignments, compared with 23% of unambiguous pairs belonging to the same classes (**Figure 2F**). Disagreement was significantly higher for flagged pairs in five of the six ambiguity types tested, with cHS|cWW the exception (**Supplementary Figure S9A**). Such ambiguity can arise when hydrogen bonds involve only atoms shared by two base edges, such as adenine N6, which belongs to both the Watson-Crick and Hoogsteen edges (see Methods).

For example, RUBRIC labels U1880-C1856 in PDB 8OVE as cWH|cHH. barnaba and RNApolis assign this pair to cWH, RNAView to cHH and FR3D to tHH (**Supplementary Figure S9B**). Similar disagreements occur for G2447-C2452 in PDB 7ABZ, labeled cWH|cWW by RUBRIC, and U2315-A2451 in PDB 8FR8, labeled tHH|tWH (**Supplementary Figure S9C and D**). By reporting candidate classes, RUBRIC makes these uncertain assignments visible to the user.

### RUBRIC scores base-pair geometry and reports individual penalties

Building on the approach for validating the bond lengths and angles of nucleic acids recommended by the Nucleic Acid Valence Geometry (NA-VAL) Working Group [63] and described in its corresponding publication by Černý et al. [22], RUBRIC evaluates each unambiguously classified base pair using ten parameters, penalizing deviations from reference geometries (see Methods). **Figure 3A** illustrates the scoring procedure for the opening angle of A-U cWW pairs. Each parameter’s contribution is reported separately, revealing the sources and severity of geometric deviations (**Figure 3C**). For example, two cWW pairs with typical geometry, G2456-C2467 in PDB 9AXU and A42-U428 in PDB 7UNW, scored 100 with no penalties (**Figure 3B**). The distorted G-C pair G11-C41 in PDB 4V2S scored 45.4, with penalties for eight of the ten parameters. The distorted A-U pair A81-U88 in PDB 7D6Z scored 73.8, with penalties for three parameters, dominated by stagger and propeller (**Figure 3B and C**).

**Figure 3:**
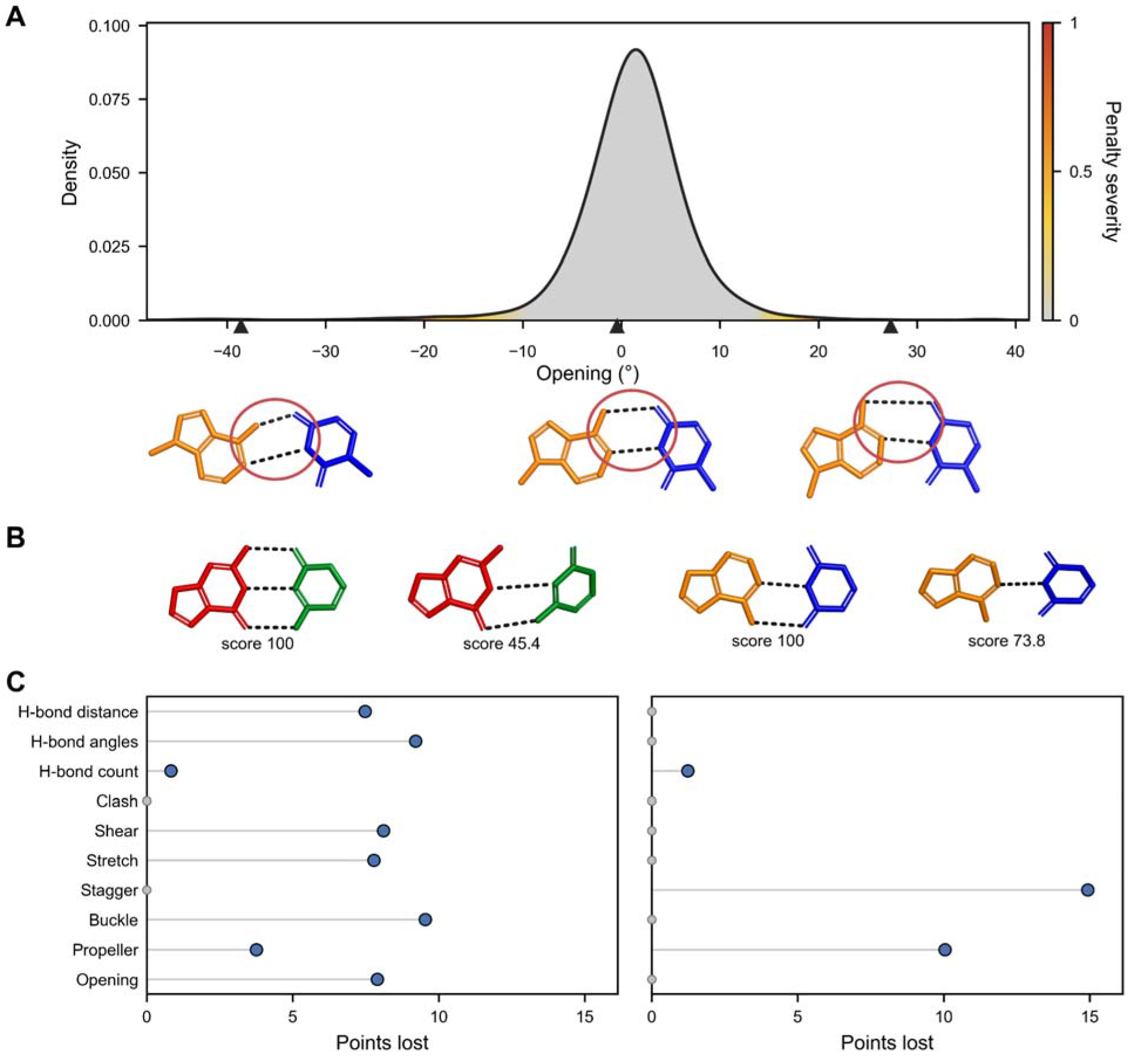
Base-pair scoring against reference geometry. (A) Reference distribution of the opening angle of A-U cWW pairs, shaded by the penalty severity assigned at each value. Triangles mark the opening of the three pairs shown below, drawn at the same scale: PDB 6Y2L, chain S2, A1293-U1306 (opening −38.6°, total score 72.5); PDB 9I78, chain L1, A2851-U2868 (−0.4°, total score 100); and PDB 6YSI, chain 1, A2780-U2794 (+27.3°, total score 72.5). Red circles mark the Watson-Crick edge of each pair. (B) G-C (left) and A-U (right) cWW pairs with ideal geometry (PDB 9AXU, chain 2, G2456-C2467; PDB 7UNW, chain A, A42-U428) and distorted geometry (PDB 4V2S, chain Q, G11-C41; PDB 7D6Z, chain f, A81-U88), with their scores. (C) Points lost to each of the ten scored parameters by the distorted G-C (left) and A-U (right) pairs in B; a small gray dot at 0 marks a parameter that costs no points. Bases are colored G red, C green, A orange and U blue; dashed lines are hydrogen bonds.

### RUBRIC base-pair scores complement standard quality measures

We next asked whether RUBRIC base-pair scores provide information beyond that captured by standard measures of experimental and model quality. We compared the mean base-pair score per structure with five quality measures: experimental resolution, clashscore, backbone suiteness, Rfree for X-ray structures and average Q-score for cryo-EM structures (**Supplementary Figure S6A-E**). Each measure explained 8-22% of the variation in base-pair scores (R²). Some association is expected because all measures except suiteness contributed to selection of the curated reference set (**Supplementary Table S2**). These modest associations suggest that base-pair scores provide complementary geometric information. Mean scores showed no detectable linear association with log-transformed RNA length (R² < 0.001; **Supplementary Figure S6F**).

About two thirds of base pairs scored 100, and scores below 80 were uncommon among both canonical and noncanonical pairs (**Supplementary Figure S10**). The average base-pair score for whole structures also differed across experimental methods, with the pattern depending on structure size (**Supplementary Figure S7**). Among structures with 100 or fewer scored pairs, median scores were lower for cryo-EM and NMR than for X-ray structures (96.7 and 97.0 versus 98.8). Among structures with 1,000 or more scored pairs, cryo-EM structures had higher median scores than X-ray structures (97.9 versus 95.9; **Supplementary Figure S7A**). Backbone suiteness differed in the same direction for the smallest and largest structures **(Supplementary Figure S7B).** With 100 or fewer scored pairs, median suiteness was lower for cryo-EM and NMR than for X-ray structures (0.46 and 0.42 versus 0.56), and with 1,000 or more scored pairs, it was higher for cryo-EM than for X-ray structures (0.51 versus 0.48). Because resolution and structure-size distributions differ across methods, these comparisons do not establish that one method produces better models than another.

### RUBRIC’s backbone flags track model quality and base-pairing context

RUBRIC assigns each backbone suite to one of the 54 conformers of the RNA Ontology Consortium or reports it as unassignable (see Methods) [12,20,64]. Because unassignable suites can reflect either modeling errors or valid conformations missing from the reference library, we examined how their frequency varies with model quality and structural context. The fraction of unassignable suites increased from 7% in structures determined at 1.8 Å or better to 27% at 3.0-3.5 Å (**Supplementary Figure S2B**). It also increased from 11% in structures with a clashscore of 2 or lower to 31% in those with a clashscore above 20 (**Supplementary Figure S2C**). These associations link assignment failures to poorer experimental resolution and model geometry, although unassignable suites persisted even at high resolution. Pucker outliers were rare in our curated high-quality reference structures, occurring in 0.05% of riboses (**Supplementary Figure S2D**).

Within the curated reference set, backbone assignments also varied with base-pairing context. Suites associated with canonical pairs were unassignable in 11% of cases, compared with 12 to 40% for noncanonical pairs, depending on the LW class (**Supplementary Figure S11A**). The A-form 1a conformer accounted for 71% of suites associated with canonical pairs, compared with 20 to 69% across noncanonical classes. In every class, unassignable suites were 1.7 to 3.6 times more frequent at C2′-endo than at C3′-endo riboses (**Supplementary Figure S11B**). These patterns suggest that noncanonical pairs and C2′-endo riboses may contain valid conformations missing from the reference library, although residual modeling errors could also contribute.

### Flagged geometry improves with higher resolution and targeted re-refinement

To test whether RUBRIC identifies correctable modeling problems, we made two comparisons: independently determined structures of the same RNA at different resolutions, and original models versus models re-refined against the same diffraction data (**Figure 4**). These comprised 162 molecules determined independently at higher resolution using the same experimental method and 1,521 X-ray structures re-refined against the original diffraction data by PDB-REDO. Base-pair score comparisons were restricted to matched pairs assigned to the same LW class in both models. PDB-REDO restrains Watson-Crick hydrogen bonds and base stacking, but not backbone torsions or ribose pucker (see Methods) [21,62]. These comparisons therefore examine changes associated with higher-resolution data and with re-refinement of unchanged data.

**Figure 4:**
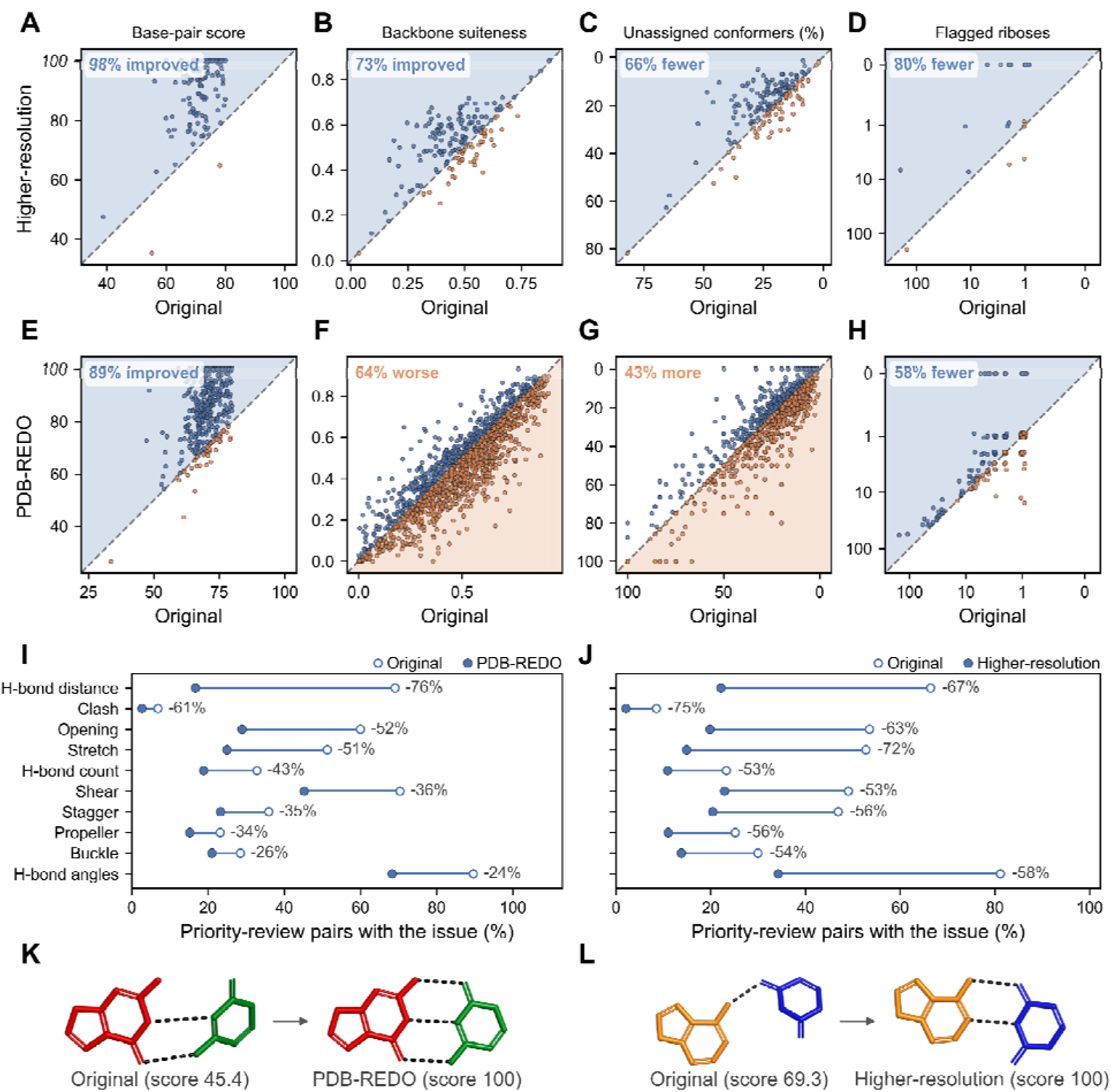
RUBRIC measures of original models compared with models at highe resolution and after PDB-REDO re-refinement. (A-D) Molecules with a second entry at higher resolution and (E-H) structures re-refined by PDB-REDO, plotted against the original model: (A and E) mean score of the base pairs in priority review (score below 80) in the original model; (B and F) backbone suiteness; (C and G) percentage of unassignable suites; (D and H) number of flagged riboses. Each point compares one original model with its re-refined or higher-resolution model; blue points improved and orange points worsened, the dashed diagonal marks no change, and the axes of C, D, G and H are inverted so that improvement lies above the diagonal. Numbers of molecules (A-D) and structures (E-H): (A) 115, (B) 162, (C) 160, (D) 30, (E) 364, (F) 1,513, (G) 1,189, (H) 192. (I and J) Percentage of originally flagged pairs showing each issue in the original model (open circles) and after re-refinement (I) or at higher resolution (J) (filled circles), with the relative change. The same cohort of originally flagged, matched pairs is followed in both models, and per-structure or per-molecule percentages are averaged across units. (K) A G-C pair (PDB 4V2S, chain Q, G11-C41) before and after PDB-REDO re-refinement. (L) A U-A pair in PDB 6V3D (chain AN1, U1167-A1175) and in PDB 7UVX, the same molecule determined at higher resolution (chain A, U1167-A1175). Bases are colored G red, C green, A orange, and U blue.

In higher-resolution models, the mean score of originally flagged pairs increased in 113 of the 115 molecules containing priority-review pairs, with the median across molecules rising from 71.8 to 93.6 (**Figure 4A**). Backbone suiteness increased in 73% of molecules, unassignable suites became less frequent in 66%, and pucker outliers decreased in 80% of the 30 molecules that initially contained them (**Figure 4B-D**).

PDB-REDO re-refinement improved mainly the Watson-Crick base-pair geometry it restrains. Mean scores of originally flagged pairs increased in 89% of the 364 structures containing them, with the median rising from 70.2 to 85.2 (**Figure 4E**). Backbone suiteness decreased in 64% of structures (median 0.52 to 0.47; **Figure 4F**), and unassignable suites became more frequent in 43% and less frequent in 28% (**Figure 4G**). Pucker outliers nevertheless decreased in 58% of the 192 structures that initially contained them (**Figure 4H**). These results concern geometric quality; RUBRIC does not assess fit to the diffraction data, which PDB-REDO also optimizes [62].

The base-pair changes further reflected the different scope of the two comparisons. After re-refinement, hydrogen-bond distance penalties showed the largest decrease in frequency, falling by 76% (**Figure 4I**). In higher-resolution models, all base-pair issue types became less frequent, with reductions of 53-75% (**Figure 4J**). Re-refinement moved 69% of flagged canonical pairs out of priority review, compared with 40% of noncanonical pairs (**Supplementary Figure S5A**); at higher resolution, the corresponding fractions were 82% and 73% (**Supplementary Figure S5B**). For example, the distorted G-C pair in **Figure 3B** changed from two detected hydrogen bonds to three after re-refinement, increasing its score from 45.4 to 100 (**Figure 4K**). A U-A pair similarly improved from 69.3 to 100 in a higher-resolution model (**Figure 4L**).

Because selecting pairs with low scores can produce apparent improvement through regression to the mean, we repeated the analysis in reverse: pairs flagged in the re-refined or higher-resolution model were followed back to the original. This comparison included 227 re-refined structures and 61 molecules with flagged pairs in both models (**Supplementary Figure S5C**). After re-refinement, the mean forward change was 9.8 points, compared with −0.6 points in reverse. For higher-resolution models, the corresponding changes were 15.9 and 10.4 points. Both forward-reverse differences were significant (*p* < 0.001), supporting improvement beyond that attributable to selection alone. Of 153 structures whose pairs all scored 100 before re-refinement, 86% retained those scores afterwards (**Supplementary Figure S5D**). These comparisons support RUBRIC’s ability to identify correctable geometric problems, with broader improvements in higher-resolution models and more selective improvements when refinement restrains the affected geometry.

### RUBRIC results can be inspected in PyMOL during model building

RUBRIC includes a PyMOL plug-in that displays its results on the structure (see Methods). Applied to chain A of the P4-P6 domain of the Tetrahymena group I intron (PDB 1GID), the plug-in colored each base pair by its score and marked the one ribose with a pucker outlier (**Supplementary Figure S12A**). The flagged pairs form a worklist that the user can step through, each with its score and penalized parameters. Superimposing the reference template of a flagged pair shows where the model departs from the expected geometry, as for the cWW pairs A235-U239 and C127-G195 (**Supplementary Figure S12B and C**). The ribose of U199 is a pucker outlier because its δ torsion (90°) indicates C3′-endo, whereas the perpendicular distance of its 3′ phosphate from the glycosidic bond line (2.3 Å; see Methods) indicates C2′-endo (**Supplementary Figure S12D**). RUBRIC can also write a PyMOL script (.pml) that reproduces the display in any PyMOL session, without the plug-in.

## Discussion

RUBRIC integrates base-pair annotation and geometric scoring with checks of backbone conformation and ribose pucker. It reports competing LW classes for ambiguous pairs and identifies the parameters lowering the scores of unambiguous pairs. Established annotation programs disagreed more often on pairs RUBRIC flagged as ambiguous. Flagged geometry generally improved in independently determined structures of the same RNA at higher resolution, while PDB-REDO re-refinement primarily improved the Watson-Crick geometry it restrains. Together, these results support reporting classification ambiguity and geometric quality separately and using local assessments to guide model inspection and correction.

The Leontis-Westhof scheme provides a common language for describing RNA base pairs. However, any scheme that divides a continuous range of geometries into discrete classes must have a mechanism for highlighting cases that do not fall into the classification system. Programs that force a single assignment will classify intermediate geometries differently because they apply different boundary conventions. Consistent with earlier comparisons, all seven programs agreed closely on canonical pairs but less on noncanonical pairs [31,35]. Because classification has no independent reference, agreement between programs measures consistency rather than accuracy [35]. The other programs disagreed significantly more often on the pairs RUBRIC flagged as ambiguous than on unambiguous pairs. Within the widely used Leontis-Westhof scheme, explicit ambiguity flags are needed to help users recognize pairs whose geometry does not justify a single class assignment, and RUBRIC provides such flags by reporting both candidate classes.

The paired model comparisons support RUBRIC’s ability to identify correctable geometric problems. Flagged features generally improved in independently determined structures of the same RNA at higher resolution. PDB-REDO re-refinement primarily improved the Watson-Crick geometry it restrains, whereas noncanonical pairs improved more frequently in the higher-resolution comparisons than after re-refinement [21]. These differences highlight opportunities for refinement procedures that address interactions they do not currently restrain. Detecting errors before deposition is particularly important because correcting released structures requires additional steps. Coordinate revisions must be initiated by the depositors or the wwPDB, while re-refinement by another group requires a separate entry and an associated peer-reviewed publication [65]. A corrected model therefore does not automatically replace the original, allowing errors to persist in structures that remain available for reuse. This has growing implications for AI methods, which use deposited RNA structures as training and evaluation data [15,66–69]. Undetected modeling errors can be learned as structural features or treated as ground truth when evaluating predictions. Validation during refinement and before deposition is therefore essential to limit the propagation of these errors into biological interpretations and computational models.

Several design choices help make RUBRIC’s scores interpretable. Type- and class-specific reference distributions allow noncanonical pairs to be assessed against their own typical geometries. Sparsely represented pairs require broader reference groups, which limits the specificity of their assessments. Reporting base-pair, backbone and ribose assessments separately preserves differences that a combined score could obscure: after PDB-REDO re-refinement, flagged base pairs generally improved even as backbone suiteness decreased in most structures. These assessments are intended primarily to guide local inspection. Like MolProbity [48], RUBRIC evaluates geometry from atomic coordinates rather than directly assessing agreement with experimental data. Local measures of model-to-data fit, including residue-level real-space R values for crystal structures and Q-scores for cryo-EM structures, therefore provide complementary evidence when evaluating a flagged feature [18,46]. Like other reference-based validation methods [20,22], RUBRIC may penalize rare but genuine geometries and may not detect systematic distortions present in its reference set. Decisions to modify flagged features should therefore consider both the experimental data and the structural context.

Backbone validation has a related limitation because the conformer library does not cover all valid RNA backbone geometries [12,64]. In the curated structures, unassignable suites were more common near noncanonical pairs and C2′-endo riboses (**Supplementary Figure S11**). These contexts may be particularly useful for extending the reference library, although unassignable suites can also reflect residual modeling errors. RUBRIC’s interaction coverage is likewise incomplete: it does not classify pairs supported only by hydrogen bonds to ribose or phosphate atoms, or assess RNA contacts with proteins and small molecules. Its flags therefore provide a starting point for inspection rather than a complete assessment of structural correctness.

RUBRIC addresses a gap in RNA structure validation by assessing canonical and noncanonical base-pair geometry together with backbone conformation and ribose pucker. Analysis of a complete human ribosome in under a second makes repeated validation practical throughout refinement. Its PyMOL interface supports inspection of flagged features, while machine-readable JSON output enables automated analysis and provides a basis for integration into refinement programs such as PHENIX [70,71]. These local assessments also support targeted curation of structural datasets, allowing researchers to identify interactions or nucleotides requiring review or exclusion while retaining usable regions of a model. Such curation is increasingly important for RNA structure-prediction methods, whose training and evaluation depend on the quality of deposited structures [15,66–69]. RUBRIC is open source (https://github.com/YesselmanLab/RUBRIC), and we welcome community contributions to extend its capabilities. To support widespread use, we provide PDB-wide RNA analyses at https://yesselmanlab.github.io/RUBRIC-PDB-live, updated weekly and accessible through a graphical interface and downloadable reports for automated analysis and machine-learning workflows. These resources make detailed geometric validation practical both during model refinement and when curating the structural data underlying biological analyses and AI development.

## Data availability

All data and software used in this study are available. All analyses were performed with RUBRIC version 1.0, available at https://github.com/YesselmanLab/RUBRIC (archived on Zenodo, https://doi.org/10.5281/zenodo.24324565). The processed data are available on Zenodo (https://doi.org/10.5281/zenodo.24214090). All code used to generate the analyses and figures is available on GitHub at https://github.com/YesselmanLabPublications/2026_Rubric_paper (archived on Zenodo, https://doi.org/10.5281/zenodo.24324880).

## Acknowledgments

We would like to thank members of the Yesselman lab for their thoughtful and helpful comments in improving this study. Computational resources were provided by the University of Nebraska’s Holland Computing Center, which is supported by the UNL Office of Research and Innovation and the Nebraska Research Initiative.

## Declaration of generative AI and AI-assisted technologies

During the preparation of this work, we used Anthropic’s Claude and OpenAI’s ChatGPT and Codex to improve the clarity of the manuscript text. Claude also assisted in implementing RUBRIC and the analysis code. All plots are generated from the underlying data by the analysis code, which is publicly deposited. We reviewed all content produced with the assistance of these tools and take full responsibility for the content of the published article.

## Contributions

J.D.Y. conceived this study. K.J.D. wrote the code and performed the analysis with assistance from J.D.Y. J.D.Y. and K.J.D. wrote the paper.

## Supplementary Information Supplementary Figures

**Figure S1:**
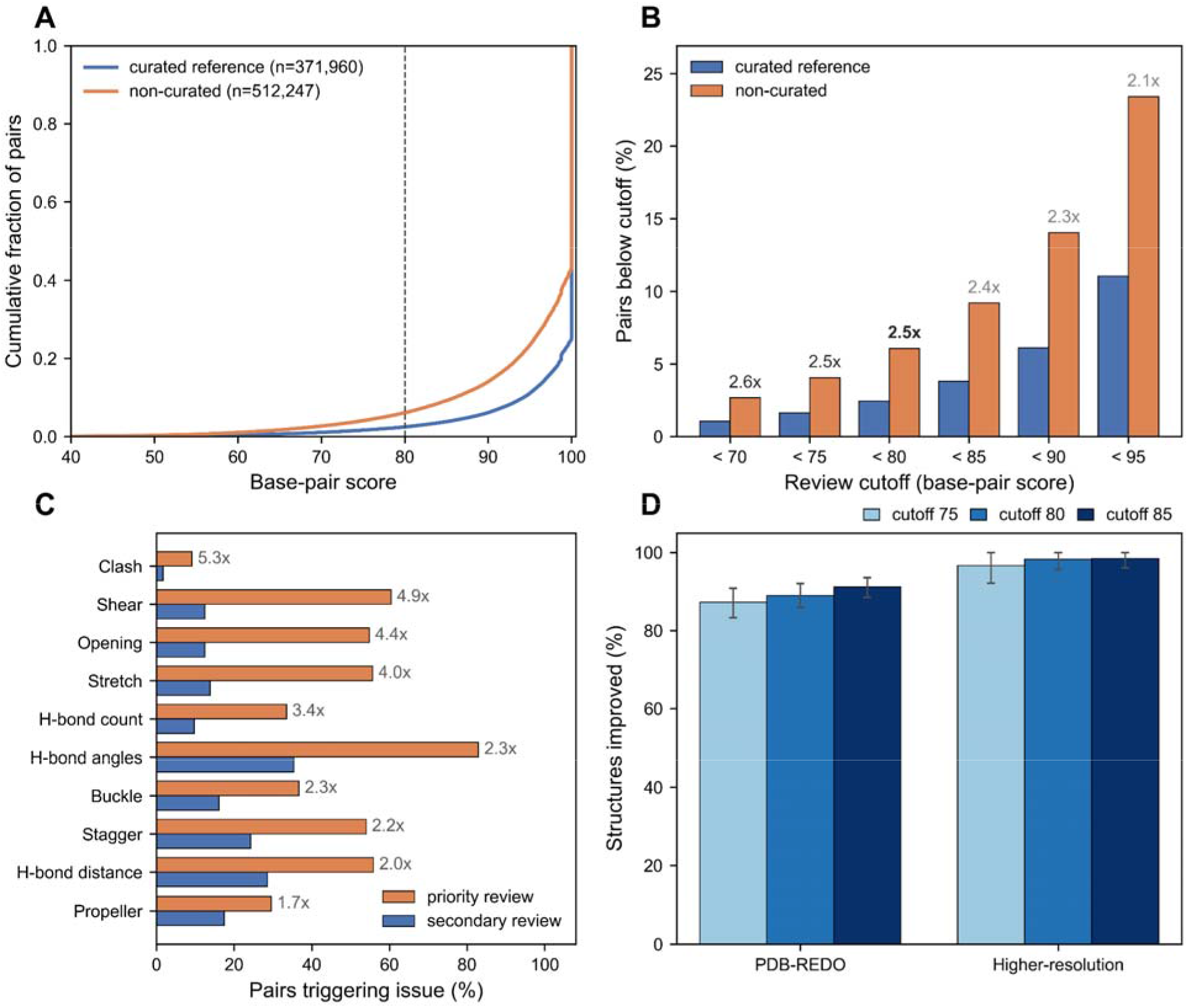
Base-pair score distributions and choice of the priority-review cutoff. (A) Cumulative distributions of base-pair scores in the curated reference set (*n = 371,960 pairs*) and the non-curated set (*n = 512,247 pairs*); the dashed line marks the cutoff of 80. (B) Percentage of pairs scoring below each cutoff in the two sets, with the fold enrichment in the non-curated set; the cutoff used is shown in bold. (C) Percentage of priority-review (score below 80) and secondary-review (80 to below 100) pairs showing each issue, with the fold enrichment in priority review. (D) Percentage of structures whose flagged pairs improve after PDB-REDO re-refinement or at higher resolution, with the review cutoff at 75, 80 and 85; error bars are 95% bootstrap confidence intervals (1,000 resamples of structures or molecules).

**Figure S2:**
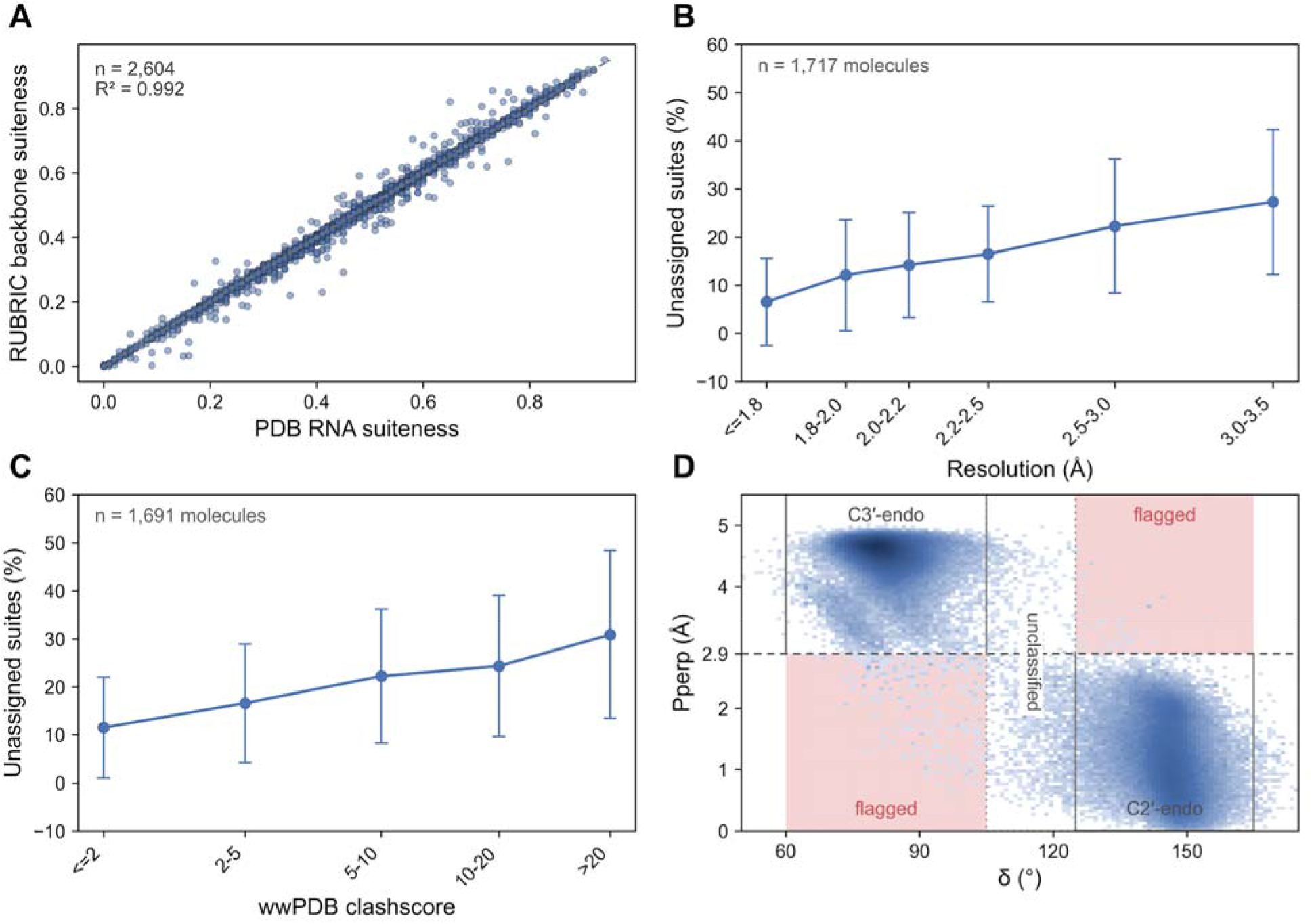
Backbone suite and ribose pucker assessment. (A) Backbone suiteness computed by RUBRIC against the RNA suiteness reported by the wwPDB (*n* = 2,604 structures). (B and C) Percentage of unassignable suites against (B) resolution and (C) wwPDB clashscore. Points are means and error bars one standard deviation of the per-molecule rates; numbers of molecules are given in each panel. (D) Ribose pucker of 855,264 residues in the curated reference structures, shown as δ against Pperp (logarithmic density). Outlined regions mark agreement between the two estimates (C3′-endo: δ 60-105° and Pperp ≥ 2.9 Å; C2′-endo: δ 125-165° and Pperp < 2.9 Å); red regions mark pucker outliers, in which the two estimates disagree; the dotted column marks δ between the two ranges, reported as unclassified.

**Figure S3:**
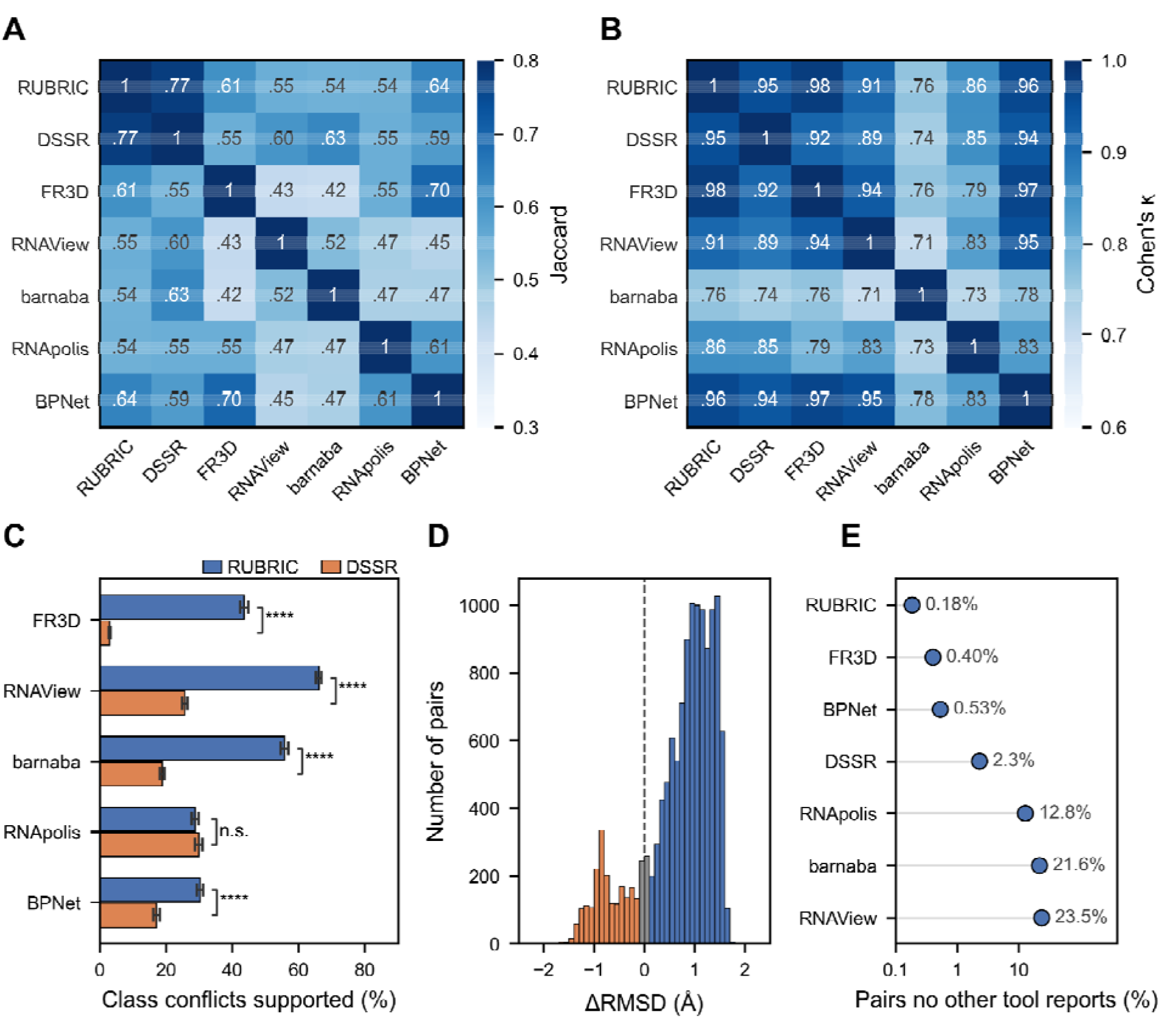
Agreement between RUBRIC and six established annotation programs. (A) Agreement in noncanonical base-pair detection between every pair of programs (Jaccard index). (B) Agreement in LW class of the noncanonical pairs detected by both programs (Cohen’s κ). (C) Pairs to which RUBRIC and DSSR assign different classes: percentage for which each of the other five programs supports the RUBRIC class (blue) or the DSSR class (orange). Error bars are 95% bootstrap confidence intervals (2,000 resamples of structures); significance is from a two-sided z-test with cluster-robust variance over structures (****, *p* < 0.0001; n.s., not significant). (D) Difference in RMSD (ΔRMSD) of the pairs in (C) to the templates of their two classes: RMSD to the template of the DSSR class minus RMSD to the template of the RUBRIC class, so that positive values (blue) are closer to the RUBRIC class, negative values (orange) closer to the DSSR class and gray bars within 0.1 Å. (E) Percentage of each program’s noncanonical pairs reported by none of the other six programs (logarithmic scale).

**Figure S4:**
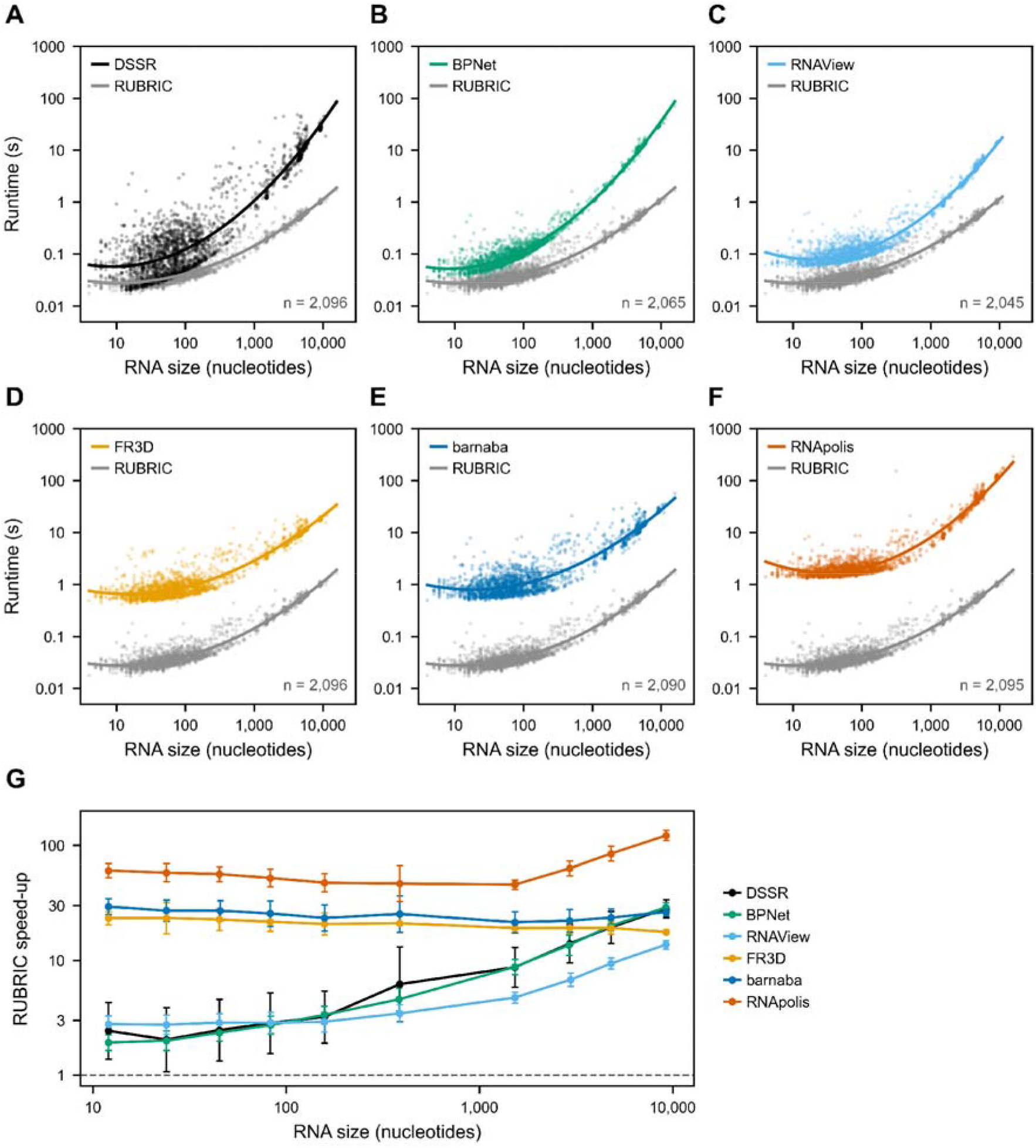
Runtime of RUBRIC and six established programs. (A-F) Runtime per structure against RNA size for (A) DSSR, (B) BPNet, (C) RNAView, (D) FR3D, (E) barnaba and (F) RNApolis Annotator, each shown with RUBRIC (gray). Lines are quadratic fits in log-log space; the number of structures is given in each panel. (G) Speed-up of RUBRIC over each program against RNA size. Points are geometric means within logarithmically spaced size bins, merged where needed so that each contains at least 30 structures, and error bars span one geometric standard deviation above and below. The dashed line marks equal speed.

**Figure S5:**
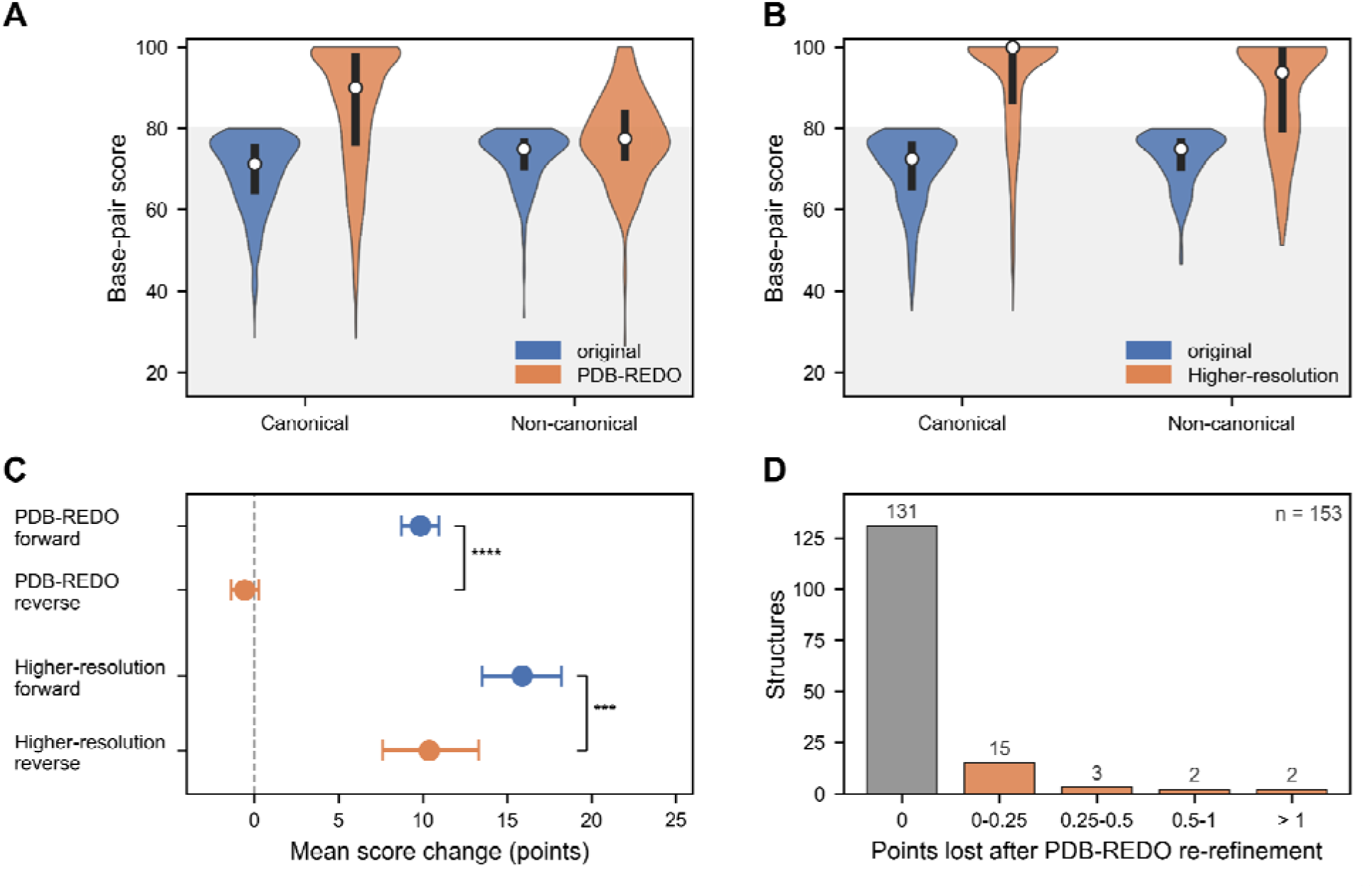
Pair-type breakdown and controls for the score change of flagged base pairs after re-refinement and at higher resolution. (A and B) Scores of canonical (A-U and G-C cWW) and noncanonical pairs that scored below 80 in the original model, before and after PDB-REDO re-refinement (A) and at higher resolution (B). White circles are medians, black bars span the interquartile range, and the gray area marks scores below 80. (C) Mean score change of pairs below 80, measured forward (selected in the original model and followed to the re-refined or higher-resolution model) and in reverse (selected in the re-refined or higher-resolution model and followed back to the original), for PDB-REDO re-refinement (227 structures) and determination at higher resolution (61 molecules); error bars are 95% bootstrap confidence intervals (4,000 resamples of structures or molecules), and significance is from a two-sided z-test on the per-structure or per-molecule difference between forward and reverse changes (***, p < 0.001; ****, p < 0.0001). (D) Points lost after PDB-REDO re-refinement by the 153 structures whose base pairs all scored 100 in the original model.

**Figure S6:**
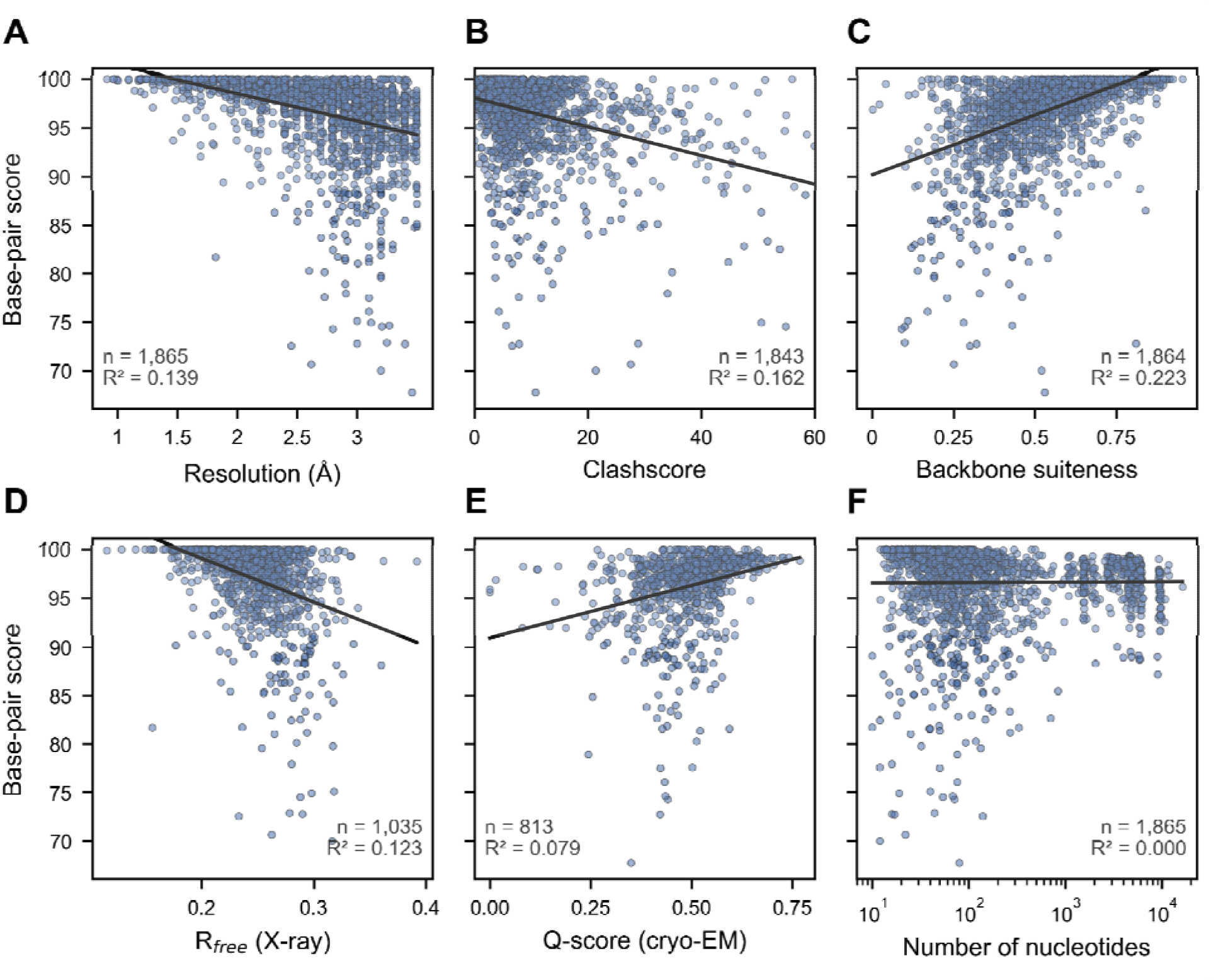
Structure-level base-pair scores against standard quality measures and size. (A) experimental resolution, (B) wwPDB clashscore, (C) RUBRIC backbone suiteness, (D) R_free_ of X-ray structures, (E) average Q-score of cryo-EM structures and (F) number of nucleotides. Each point is one structure with at least five scored base pairs. Lines are least-squares fits; the number of structures (n) and the variance explained (R²) are given in each panel.

**Figure S7:**
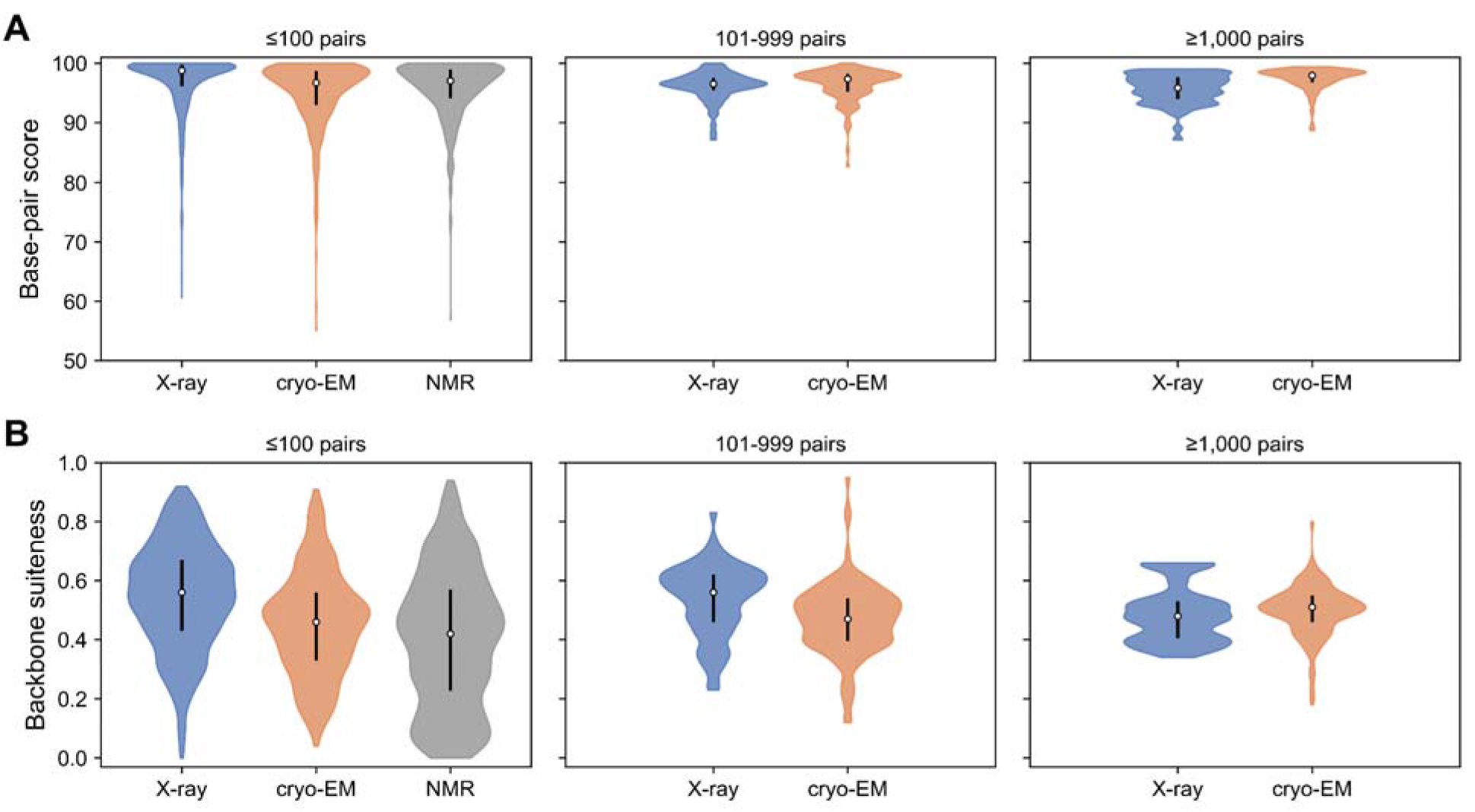
Structure-level scores by experimental method and structure size. (A) Structure-level base-pair score and (B) backbone suiteness of X-ray, cryo-EM and NMR structures, grouped by the number of scored base pairs per structure (100 or fewer, 101-999, and 1,000 or more). Violins show the distribution over structures, white circles are medians and black bars span the interquartile range. Methods with fewer than five structures in a group are not shown. NMR structures are a non-redundant set scored on the first model.

**Figure S8:**
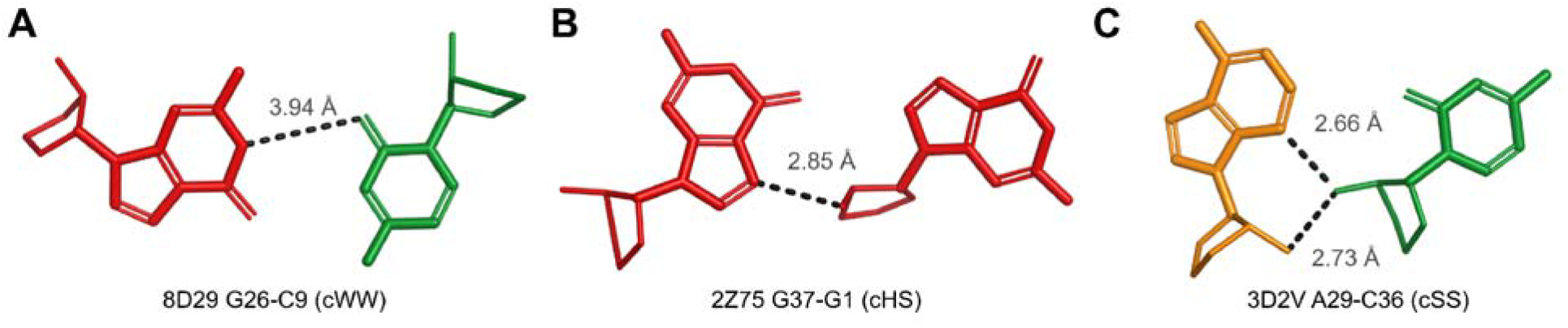
Base pairs reported by DSSR but not by RUBRIC. (A) G26-C9 in PDB 8D29 (chain R), assigned to cWW by DSSR; (B) G37-G1 in PDB 2Z75 (chains B and A), assigned to cHS (C) A29-C36 in PDB 3D2V (chain A), assigned to cSS. Dashed lines are the hydrogen bonds reported by DSSR, labeled with their donor-acceptor distances. Bases and riboses are colored G red, C green, and A orange. Each ribose drawn as thin sticks takes the color of its base.

**Figure S9:**
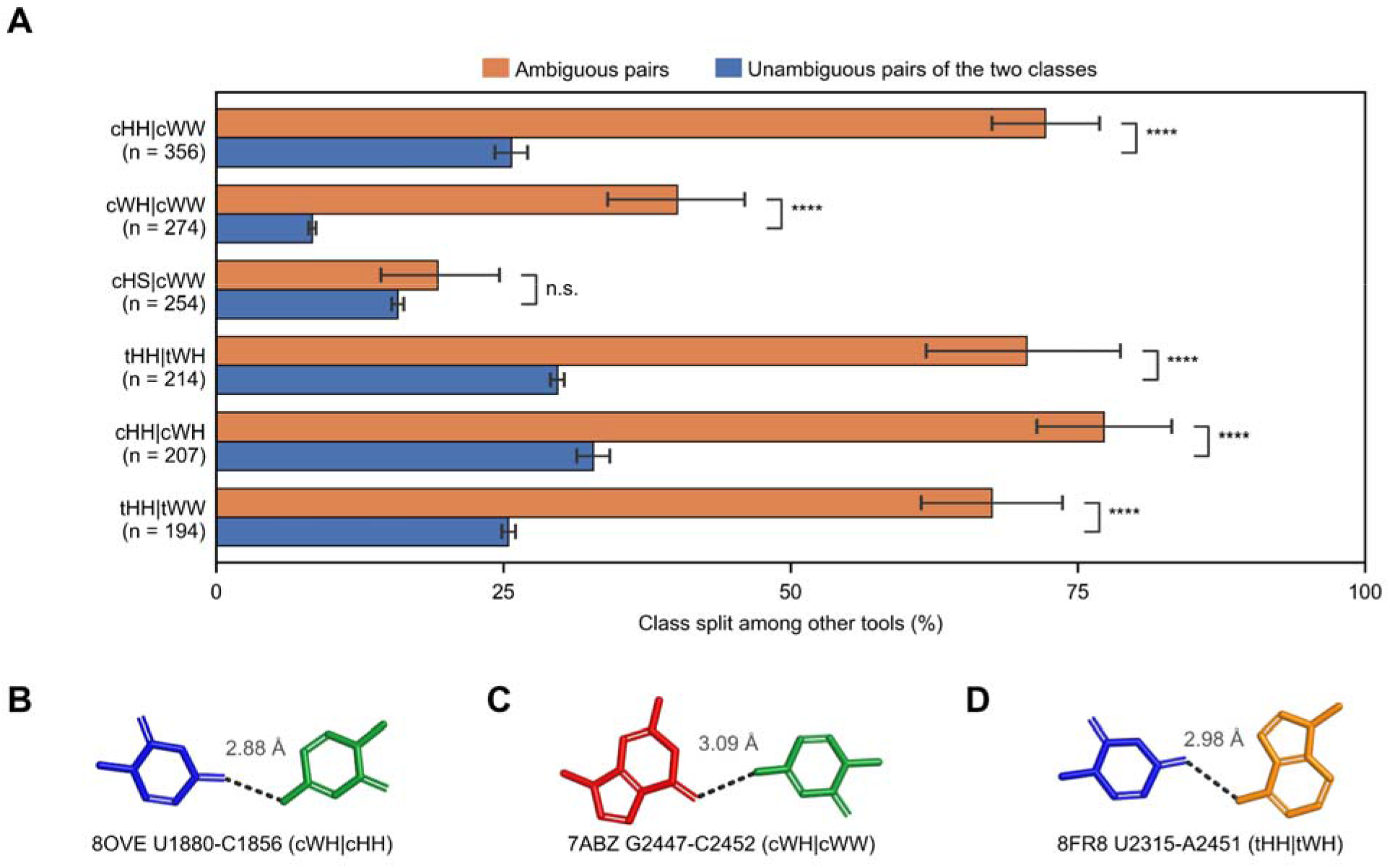
Analysis of ambiguous base pairs. (A) Percentage of RUBRIC pairs on which the other programs assign more than one LW class, for each of the six commonest ambiguity types: pairs that RUBRIC reports as ambiguous (orange) and unambiguous pairs of the two classes involved (blue). n is the number of ambiguous pairs. Error bars are 95% bootstrap confidence intervals (2,000 resamples of structures); significance is from a two-sided z-test with cluster-robust variance over structures (****, p < 0.0001; n.s., not significant). (B-D) Pairs that RUBRIC reports as ambiguous: (B) U1880-C1856 in PDB 8OVE (chain AA), cWH|cHH; (C) G2447-C2452 in PDB 7ABZ (chain 1), cWH|cWW; (D) U2315-A2451 in PDB 8FR8 (chain A), tHH|tWH. Dashed lines are hydrogen bonds, labeled with their donor-acceptor distances. Bases are colored G red, C green, A orange, and U blue.

**Figure S10:**
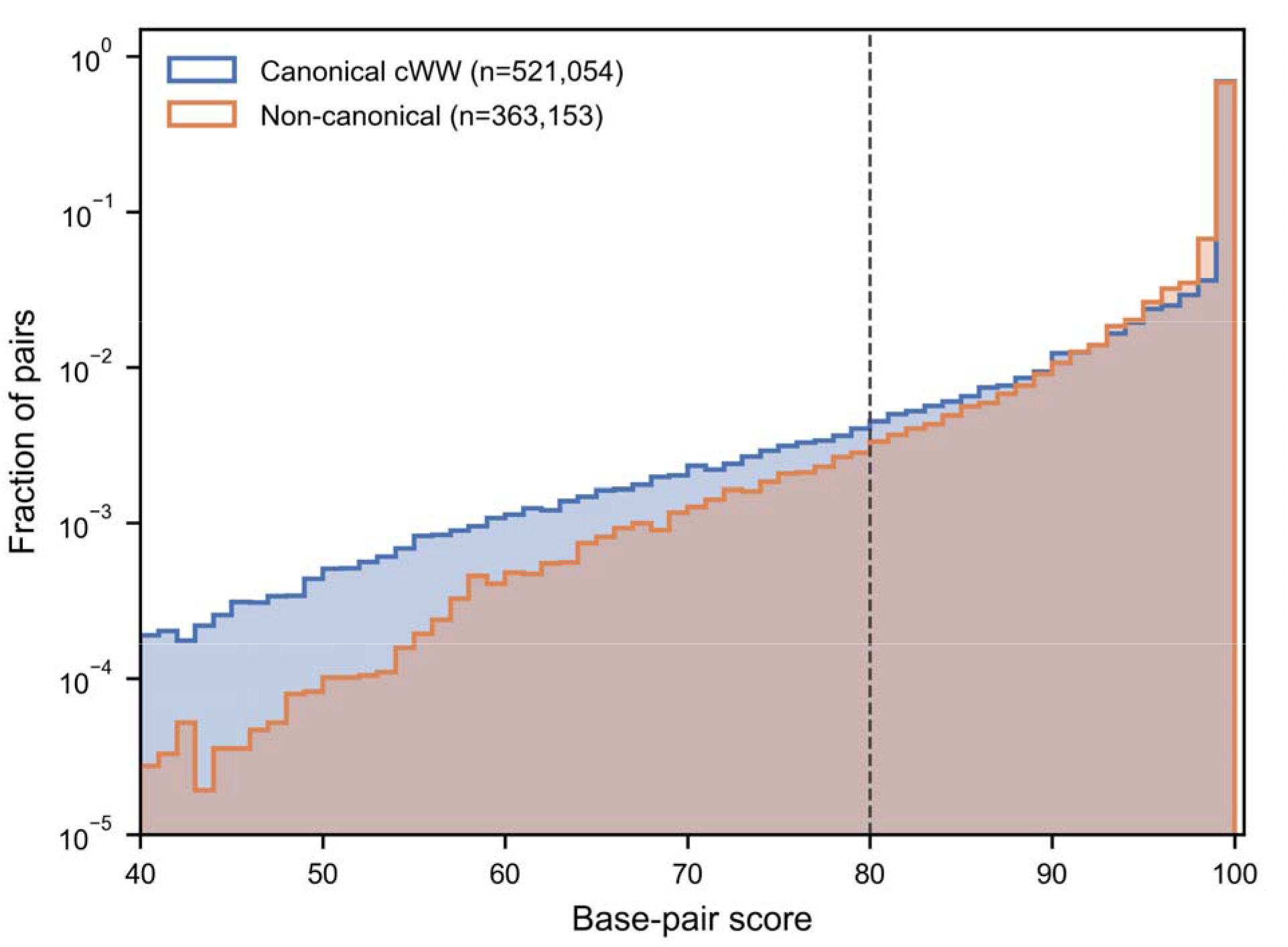
Score distributions of canonical and noncanonical base pairs. Histograms of the scores of canonical A-U and G-C cWW pairs (blue) and noncanonical pairs (orange), for the unambiguous base pairs of the non-redundant set. Each histogram shows the fraction of pairs in its group at each score. The vertical axis is logarithmic. The dashed line marks the priority-review cutoff of 80.

**Figure S11:**
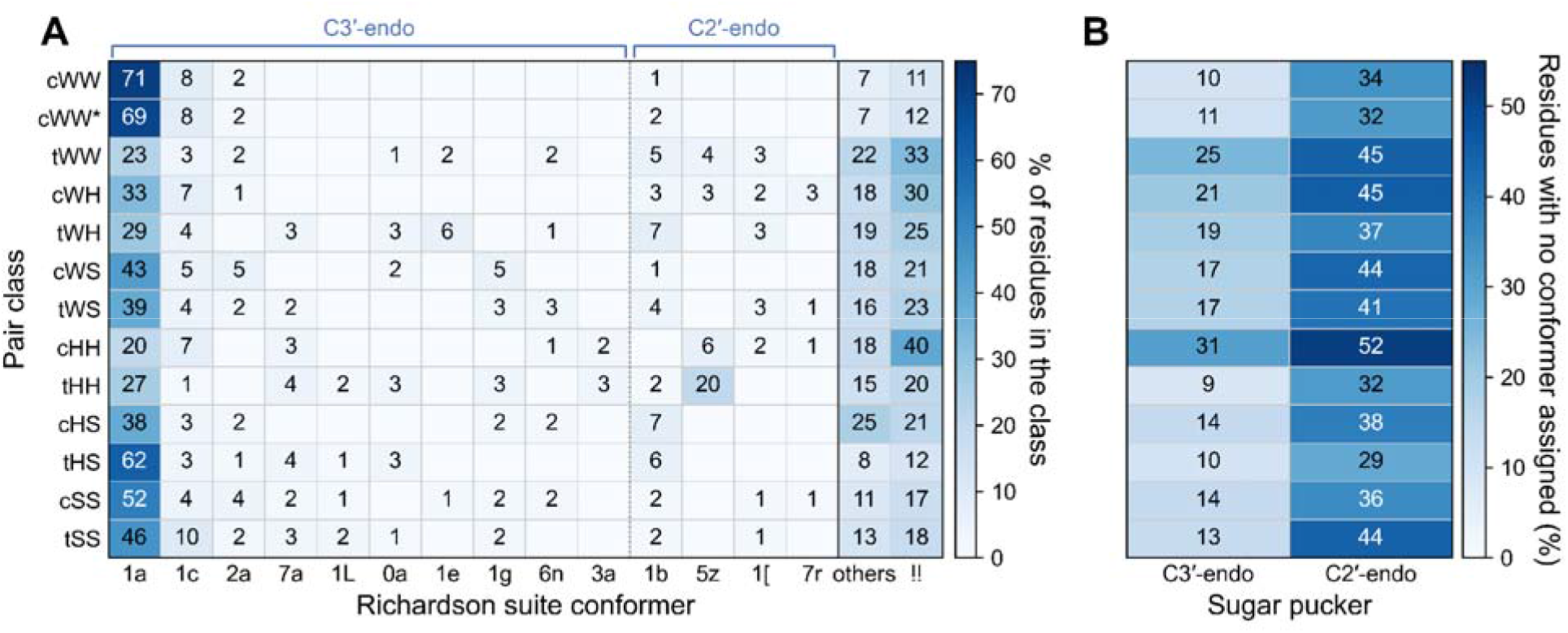
Backbone conformers and unassignable suites by base-pair class and ribose pucker. (A) Usage of Richardson suite conformers by the residues of each base-pair class, as a percentage of the residues in the class. Conformers are grouped by the pucker of their ribose (C3′-endo or C2′-endo), conformers used by less than 1% of a class are pooled as others, and !! denotes unassignable suites; cWW* denotes cWW pairs with a noncanonical sequence. (B) Percentage of residues with an unassignable suite among C3′-endo and C2′-endo riboses of each class.

**Figure S12:**
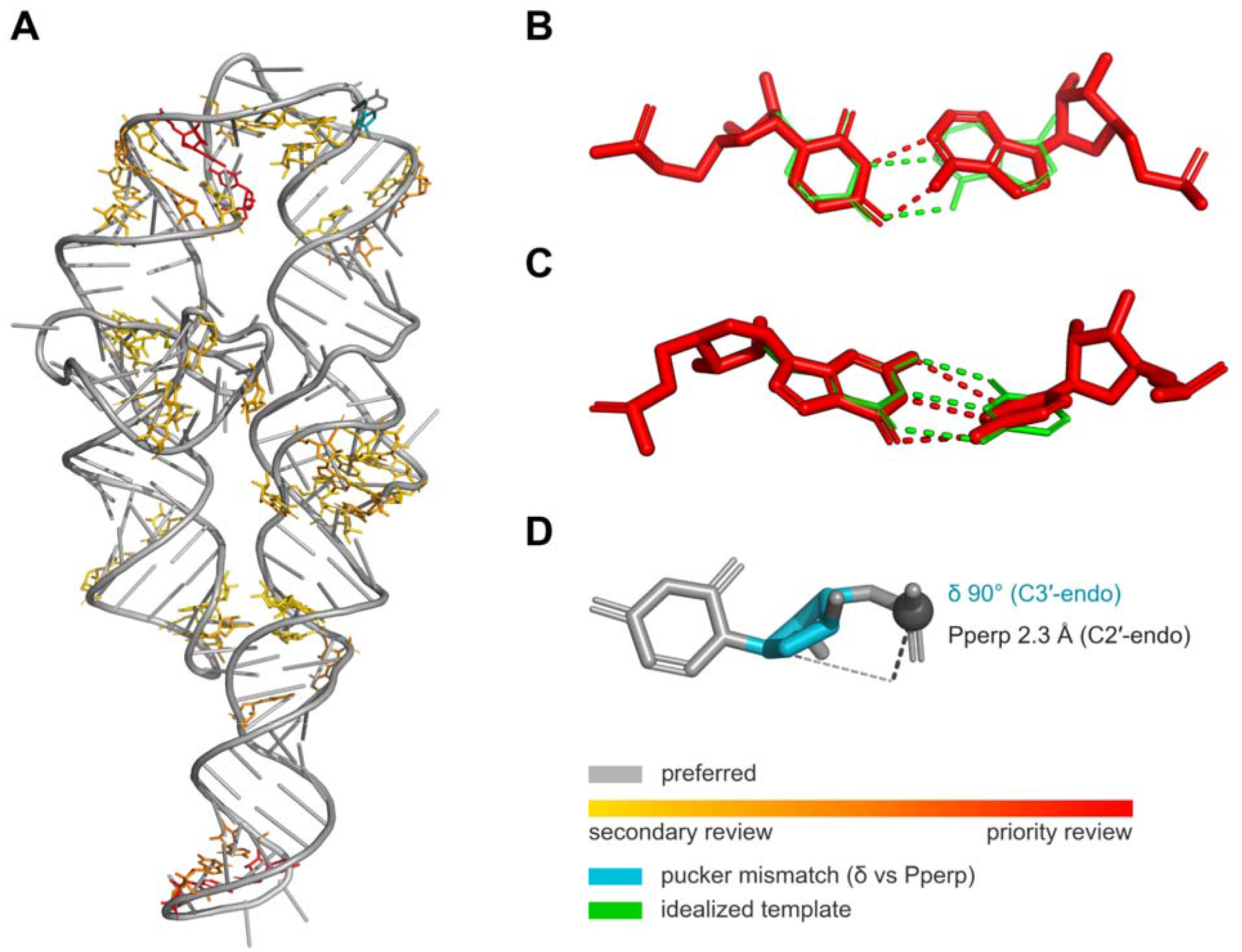
Inspection of RUBRIC results in PyMOL. (A) Chain A of the P4-P6 domain of the Tetrahymena group I intron (PDB 1GID) displayed with the RUBRIC PyMOL plug-in, with each base pair colored by its score: gray, preferred (score 100); yellow to orange, secondary review (80 to below 100); red, priority review (below 80). The ribose flagged for a pucker outlier is shown in cyan. (B) The A235-U239 cWW pair (score 74.5; H-bond distance, opening, stretch and H-bond angles flagged) and (C) the C127-G195 cWW pair (score 76.6; buckle, H-bond angles, propeller, H-bond distance and stagger flagged), each superimposed on its reference template (green). Dashed lines are hydrogen bonds of the model (red) and of the template (green). (D) The ribose of U199, whose δ torsion (90°) indicates C3′-endo whereas the perpendicular distance of its 3′ phosphate from the glycosidic bond line (Pperp, 2.3 Å) indicates C2′-endo.

## Supplementary Tables

**Table S1:**
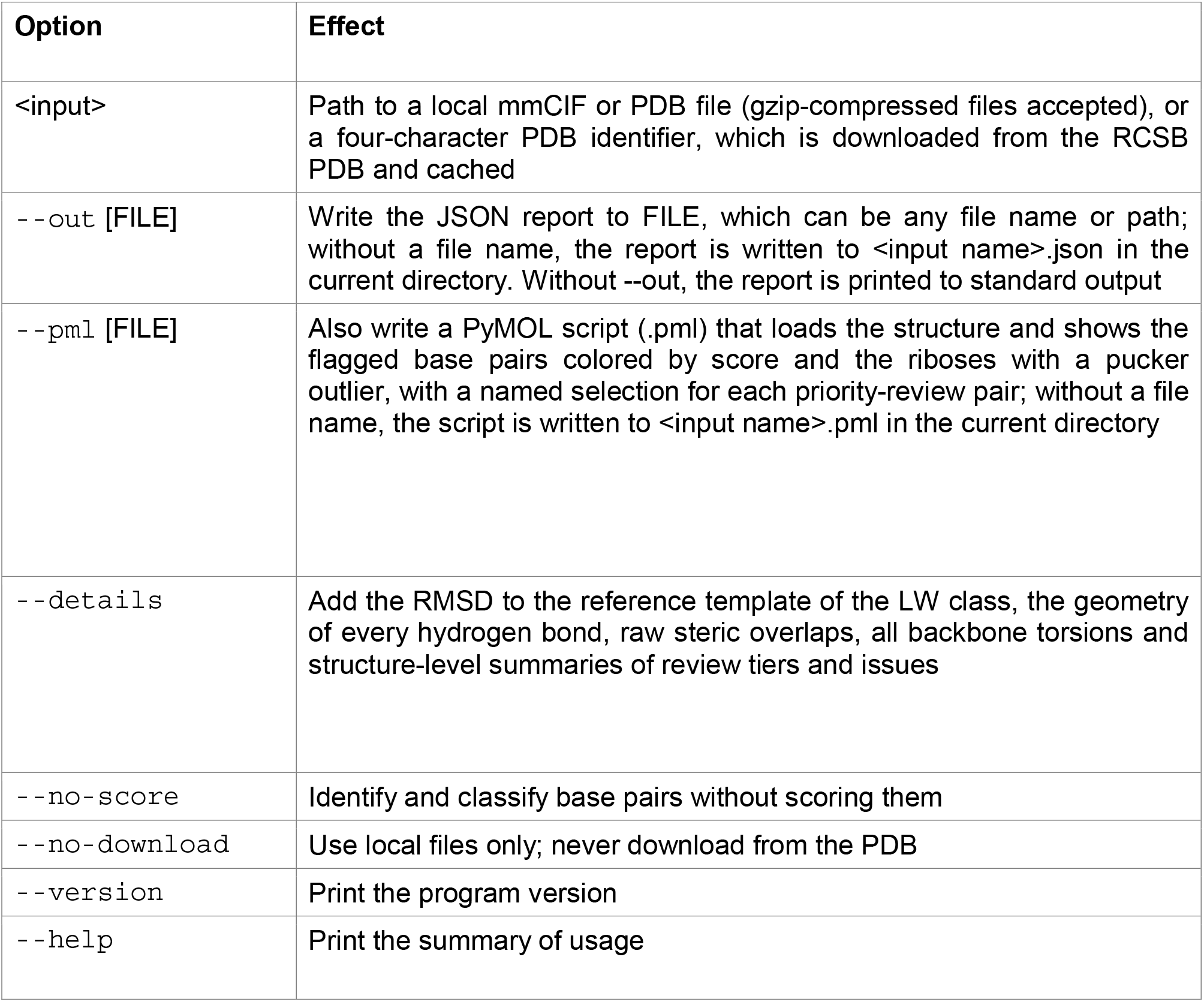
Command-line options of RUBRIC. Each option of the rubric command and what it does. Options can be given in any order after the input file.

**Table S2:** Criteria for the curated reference set. Quality criteria from the wwPDB validation reports that an entry of the non-redundant set had to meet to enter the curated reference set. An entry was included in the curated set only if it met every criterion for its experimental method.

| **Applies to** | **Criterion** |
|---|---|
| All entries | Member of the non-redundant set; ≥ 1 nucleotide |
| All entries | Bond-length RMSZ ≤ 2.0 and bond-angle RMSZ ≤ 2.0 |
| All entries | Clashscore ≤ 20 |
| X-ray | Resolution ≤ 3.0 Å; R<sub>free</sub> ≤ 0.28; R<sub>free</sub> − R<sub>work</sub> ≤ 0.05 |
| Cryo-EM | Reported resolution ≤ 3.0 Å; average Q-score ≥ 0.5 |

**Table S3:** Penalty weights of the ten scored parameters. Mean severity of each parameter over the scorable base pairs of the curated (371,960 pairs) and non-curated (512,247 pairs) reference sets, the natural logarithm of their ratio, and the resulting weight, rescaled so that the ten weights sum to 100. Parameters are listed by decreasing weight.

| **Parameter** | **Mean severity, curated** | **Mean severity, non-curated** | **ln(ratio)** | **Weight** |
|---|---|---|---|---|
| Stagger | 0.0194 | 0.0556 | 1.053 | 15.05 |
| Steric clash | 0.0019 | 0.0045 | 0.867 | 12.40 |
| H-bond angles | 0.0397 | 0.0869 | 0.785 | 11.22 |
| Shear | 0.0181 | 0.0378 | 0.738 | 10.55 |
| Propeller | 0.0143 | 0.0289 | 0.702 | 10.04 |
| Buckle | 0.0158 | 0.0311 | 0.678 | 9.69 |
| H-bond distance | 0.0303 | 0.0593 | 0.670 | 9.59 |
| Stretch | 0.0189 | 0.0368 | 0.668 | 9.55 |
| Opening | 0.0166 | 0.0320 | 0.658 | 9.42 |
| H-bond count | 0.0199 | 0.0237 | 0.173 | 2.48 |

**Table S4:**
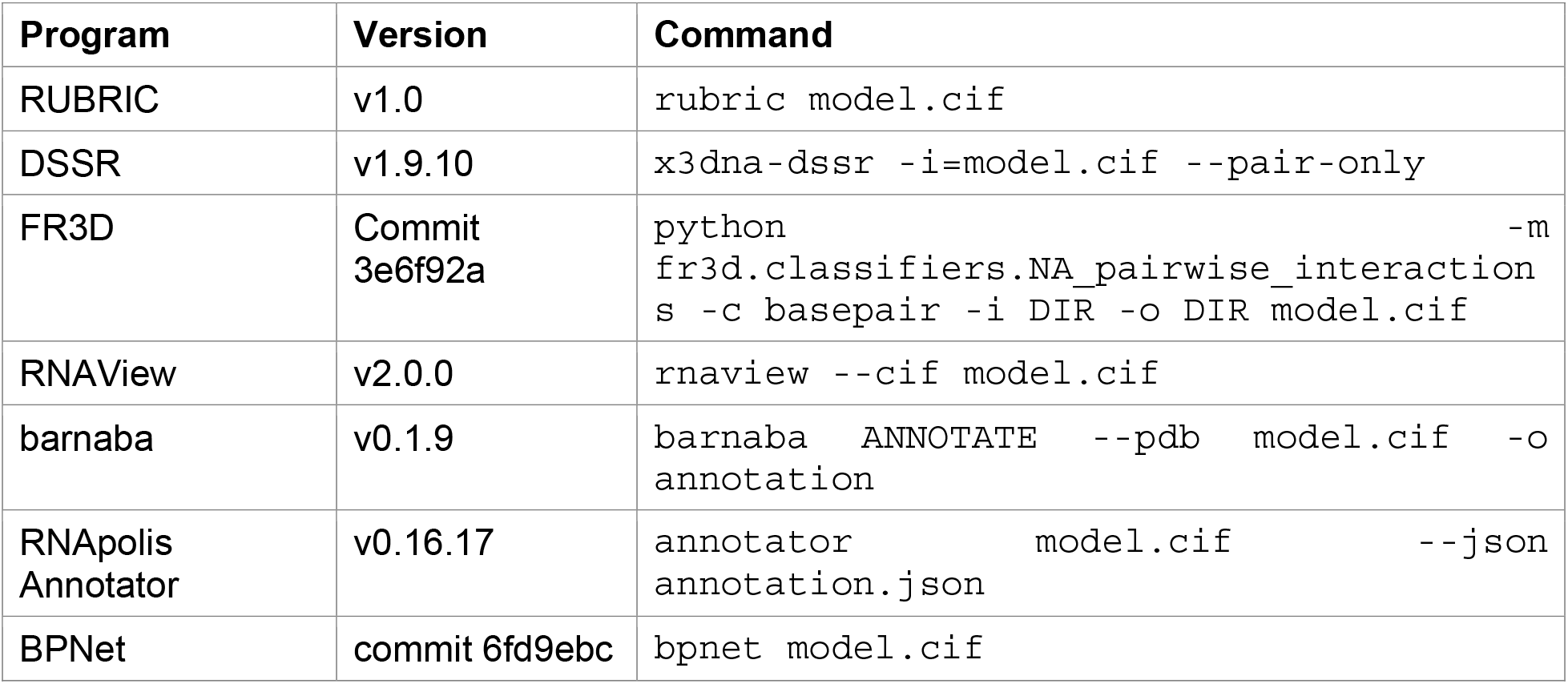
Program versions and commands. Versions and commands of RUBRIC and the six annotation programs used in the runtime benchmark and the base-pair comparison. model.cif is the input mmCIF file and DIR a working directory.

| Program | Version | Command |
|---|---|---|
| RUBRIC | v1.0 | rubric model.cif |
| DSSR | v1.9.10 | x3dna-dssr -i=model.cif --pair-only |
| FR3D | Commit 3e6f92a | python -m fr3d.classifiers.NA_pairwise_interactions -c basepair -i DIR -o DIR model.cif |
| RNAView | v2.0.0 | rnaview --cif model.cif |
| barnaba | v0.1.9 | barnaba ANNOTATE --pdb model.cif -o annotation |
| RNApolis Annotator | v0.16.17 | annotator model.cif --json annotation.json |
| BPNet | commit 6fd9ebc | bpnet model.cif |

**Table S5:** Detection agreement with selected and classified pairs. Detection agreement (Jaccard index) between RUBRIC and each program on 2,660 structures, computed with RUBRIC’s selected pairs or with all pairs it classified, whether selected or not.

| **Program** | **Canonical, selected** | **Canonical, all classified** | **Non-canonical, selected** | **Non-canonical, all classified** |
|---|---|---|---|---|
| DSSR | 0.980 | 0.981 | 0.773 | 0.836 |
| FR3D | 0.962 | 0.938 | 0.613 | 0.546 |
| RNAView | 0.977 | 0.968 | 0.549 | 0.575 |
| barnaba | 0.981 | 0.981 | 0.543 | 0.608 |
| RNApolis | 0.976 | 0.962 | 0.545 | 0.521 |
| BPNet | 0.975 | 0.954 | 0.638 | 0.582 |

## Supplementary Sample Output

Excerpt of the JSON report written by RUBRIC for PDB 1GID with default options. The structure-level summary is followed by one priority-review base pair (A235-U239; **Supplementary Figure S12B**), one backbone residue with suggested torsion corrections and one ribose with a pucker outlier (U199; **Supplementary Figure S12D**). Only one entry of each list is shown, and the reference frames of the base pair are omitted. Within issue_details, tier is the classification of each penalized value by Černý et al. [22] Running RUBRIC with --details adds, for each base pair, the geometry of every hydrogen bond (donor and acceptor atoms, distance and angles), and torsions for every backbone suite (**Supplementary Table S1**); for the pair shown, these include an N6-O4 hydrogen bond of 2.31 Å, the source of its distance penalty.

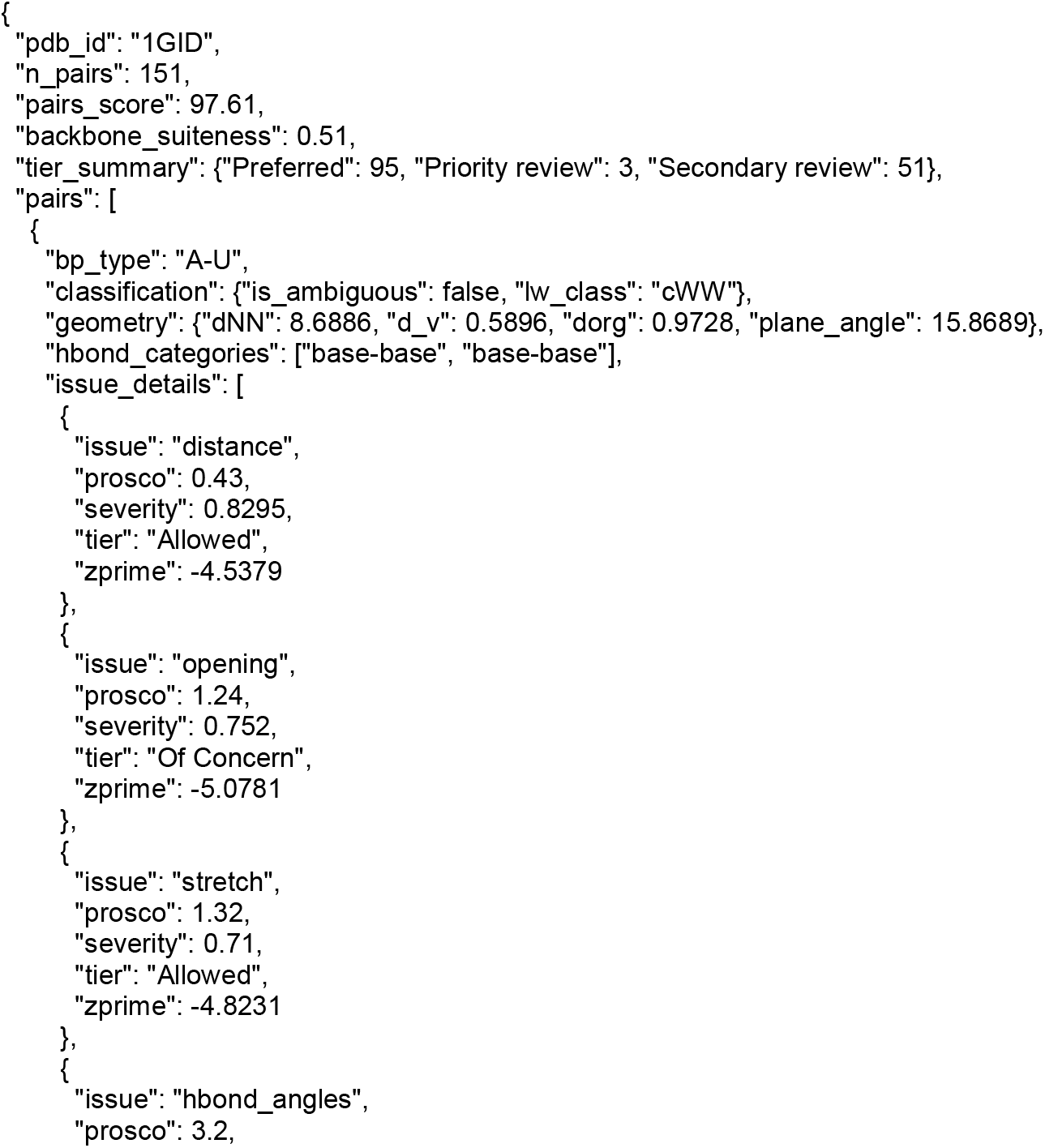
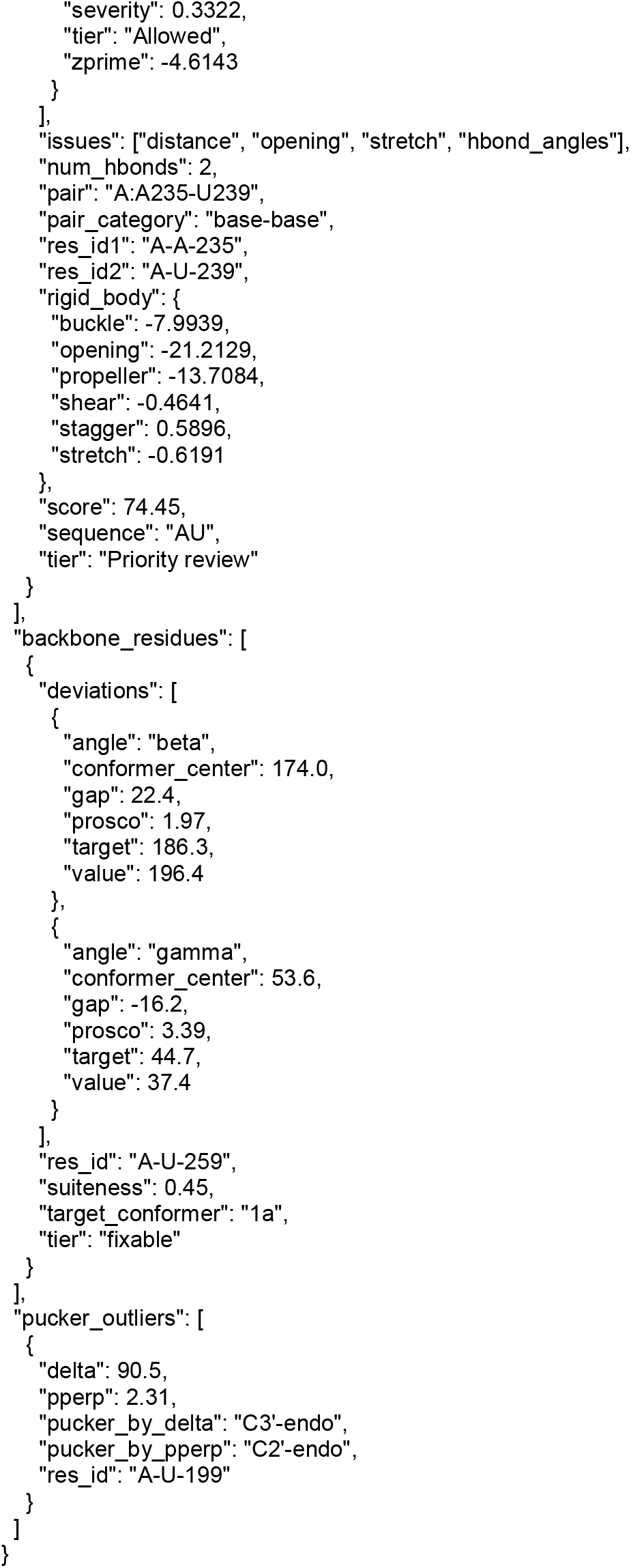

